# The *App*^*NL-G-F*^ mouse model of Alzheimer’s disease is refractory to regulatory T cell treatment

**DOI:** 10.1101/2022.03.11.483903

**Authors:** Lidia Yshii, Loriana Mascali, Lubna Kouser, Pierre Lemaitre, Marika Marino, James Dooley, Oliver Burton, Jeason Haughton, Zsuzsanna Callaerts-Vegh, Bart De Strooper, Matthew G. Holt, Emanuela Pasciuto, Adrian Liston

**Affiliations:** VIB Center for Brain and Disease Research, Leuven 3000, Belgium; KU Leuven, Department of Microbiology, Immunology and Transplantation, Leuven 3000, Belgium; KU Leuven - Department of Neurosciences, Leuven 3000, Belgium; Immunology Programme, The Babraham Institute, Babraham Research Campus, Cambridge, CB22 3AT United Kingdom; KU Leuven, Faculty of Psychology, Laboratory of Biological Psychology, 3000 Leuven, Belgium; Instituto de Investigaçāo e Inovaçāo em Saúde (i3S), University of Porto, 4200-135 Porto, Portugal

**Author notes:** equal contribution (Matthew G. Holt, Emanuela Pasciuto, Adrian Liston).

**Keywords:** Alzheimer’s Disease, APP, mouse model, regulatory T cells, IL2

## Abstract

**Background:** Alzheimer’s Disease is a neurodegenerative disease with a neuroinflammatory component. Due to the multifunctional capacity of regulatory T cells to prevent and reverse inflammation, regulatory T cells have been proposed as a potential therapeutic in Alzheimer’s Disease, either as a direct cell therapy or through the use of IL2 as a biologic to expand the endogenous population.

**Methods:** Here we characterize the longitudinal immunological changes occurring in T cells in the *App*^*NL-G-F*^ mouse model of Alzheimer’s disease.

**Results:** Age-dependent immunological changes, in both the brain and periphery, occurred in parallel in both *App*^*NL-G-F*^ mice and control *App*^*NL*^ mice. As the endogenous IL2 axis was disturbed with age, we sought to determine the effect of IL2 supplementation on disease progression. Using a genetic model of IL2 provision in the periphery or in the brain, we found that expanding regulatory T cells in either location was unable to alter the progression of key pathological events or behavioral changes.

**Conclusions:** These results suggest that either the *App*^*NL-G-F*^ mouse model does not recapitulate key regulatory T cell-dependent process of Alzheimer’s disease, or that regulatory T cell therapy is not a promising candidate for APP-mutation-driven Alzheimer’s disease.

## Background

Alzheimer’s disease (AD) is the most common neurodegenerative disease and is characterized by β-amyloid peptide (Aβ) plaques and tau tangles, complemented by astrogliosis and activated microglia [1]. However, growing evidence suggests that the amyloid cascade alone cannot recapitulate the pathogenesis of AD, indicating the involvement of other pathological processes [2]. In particular, in AD patients, T cell activation is observed in both the peripheral blood [3] and cerebral spinal fluid (CSF) [4], and T cell infiltration into the brain has been observed[5]. Moreover, AD risk variants are associated with both innate immune functions, and variants in HLA, responsible for antigen presentation to T cells [6, 7]. Further linkage to the adaptive immune system is suggested through the report of lower IL2 levels in the AD brain [8]. Taken together, inflammation is likely a fundamental player during AD progression, with infiltrating T cells a likely mediator of pathology [9].

With a potential pathophysiologic function for T cells, harnessing regulatory T cells (Tregs) has been proposed as a potential therapeutic approach. Tregs are multifunctional anti-inflammatory cells, capable of suppressing inappropriate immune activation[10]. The vast majority of research on Tregs has occurred in the peripheral context, where they increase the threshold for activation of conventional T cells. In the neurological context, this potentially allows Tregs to prevent the priming of neurodestructive T cells[11, 12]. Recently, there has been a growing appreciation that a small population of Tregs are resident in the brain tissue of mice and humans, even in the healthy context [13]. These cells can produce neurotrophic factors, such as brain-derived neurotrophic factor (BDNF) [14] and anti-inflammatory factors, such as amphiregulin, suppressing neurotoxic astrogliosis [15]. We recently demonstrated that expansion of the brain-resident Treg population, through the local provision of the key survival factor interleukin 2 (IL2) was neuroprotective in mouse models of traumatic brain injury, stroke and multiple sclerosis[16]. Together, these findings suggest that brain Tregs can play a protective role in central nervous system-related inflammation.

We considered therefore Tregs as a potential therapeutic target in AD, either through direct cell therapy or through the supplementation of the endogenous population with the survival factor IL2 [17]. The role of Tregs in AD is, however, highly contested, with different studies proposing either a protective effect of Tregs [18], a detrimental effect of Tregs [19, 20] or no effect of Tregs [21] on plaque load. Increased numbers of Tregs have been correlated with reduced microglia activation and production of inflammatory cytokines [18], but the effect of Treg interaction on phagocytic capacity of microglia is debated [20, 21]. The beneficial effect of Tregs on the cognitive performance of AD mouse model is, however, complementary [8, 18, 21]. Confounding these studies is a potential divergence in the net effect of peripheral or brain-resident Tregs [19], with most studies using a methodology that does not allow for discrimination. In order to independently test the potential benefit of therapeutic IL2-mediated Treg expansion, and to distinguish between the roles of peripheral and brain resident Tregs, here we used two specific methods for the genetic delivery of IL2 expression allowing the systemic-or brain-specific expansion of Tregs in the *App*^*NL-G-F*^ mouse model of Alzheimer’s Disease. Expansion of the Treg population was observed as expected: however, no substantial amelioration of plaque formation or cognitive decline was observed. These results suggest the pathology of the *App*^*NL-G-F*^ mouse model is refractory to Treg treatment, either through systemic or brain-directed expansion.

## Results

### Cumulative age-dependent changes in the peripheral and brain-resident immunological compartments in AD mice

Growing evidence suggests T cell involvement in AD pathogenesis [9]. We performed in depth phenotyping by high dimensional flow cytometry of the T cell compartment in the *App*^*NL-G-F*^ and in the control *App*^*NL*^ strain, and at different stages of disease progression. Specifically, before plaques deposition (2 months), after plaques deposition (4 months), and when cognitive impairment is detectable (9 months) in the *App*^*NL-G-F*^ [22]. We first assessed the peripheral compartment, represented by the spleen, blood and the draining cervical lymph nodes (cLN). Classification of T cell populations by tSNE analysis (**Supplementary Figure 1**), found a decline in naïve CD4 T cells with age across spleen and blood, consistent in both the *App*^*NL*^ and *App*^*NL-G-F*^ mice (**Figure 1A**). Naïve CD4 T cells were displaced by increased CD8 T cells, and, in some mice, large increases in memory CD4 T cells (**Figure 1A**). We next investigated the expression of key activation markers (**Figure 1B**) and cytokine production (**Figure 1C**), in each of the conventional CD4 T cell (**Supplementary Figure 2**), Treg (**Supplementary Figure 3**) and CD8 T cell (**Supplementary Figure 4**) lineages. While most phenotypes remained stable with age, several consistent changes observed, including a reduction in CD25 expression in Tregs (**Figure 1B, D**), an increase in amphiregulin production by Tregs (**Figure 1C, D**) and elevated TNF production by CD8 T cells, with a peak at 4 months on *App*^*NL-G-F*^ mice (**Figure 1C,D**). Changes were largely mirrored between *App*^*NL*^ and *App*^*NL-G-F*^ mice; however, cytokine production was inconsistent between the two strains, with *App*^*NL-G-F*^ mice demonstrating increased IL2 production by CD4 T cells and TNF production by CD8 T cells in the cLN at four months of age, and a slight decrease in IL2 production by CD4 T cells at 9 months in the spleen (**Figure 1D**). These results suggest that *App*^*NL-G-F*^ mice undergo predictable changes in the T cell compartment, driven largely by age rather than the that *App*^*NL-G-F*^ mutations.

**Figure 1.**
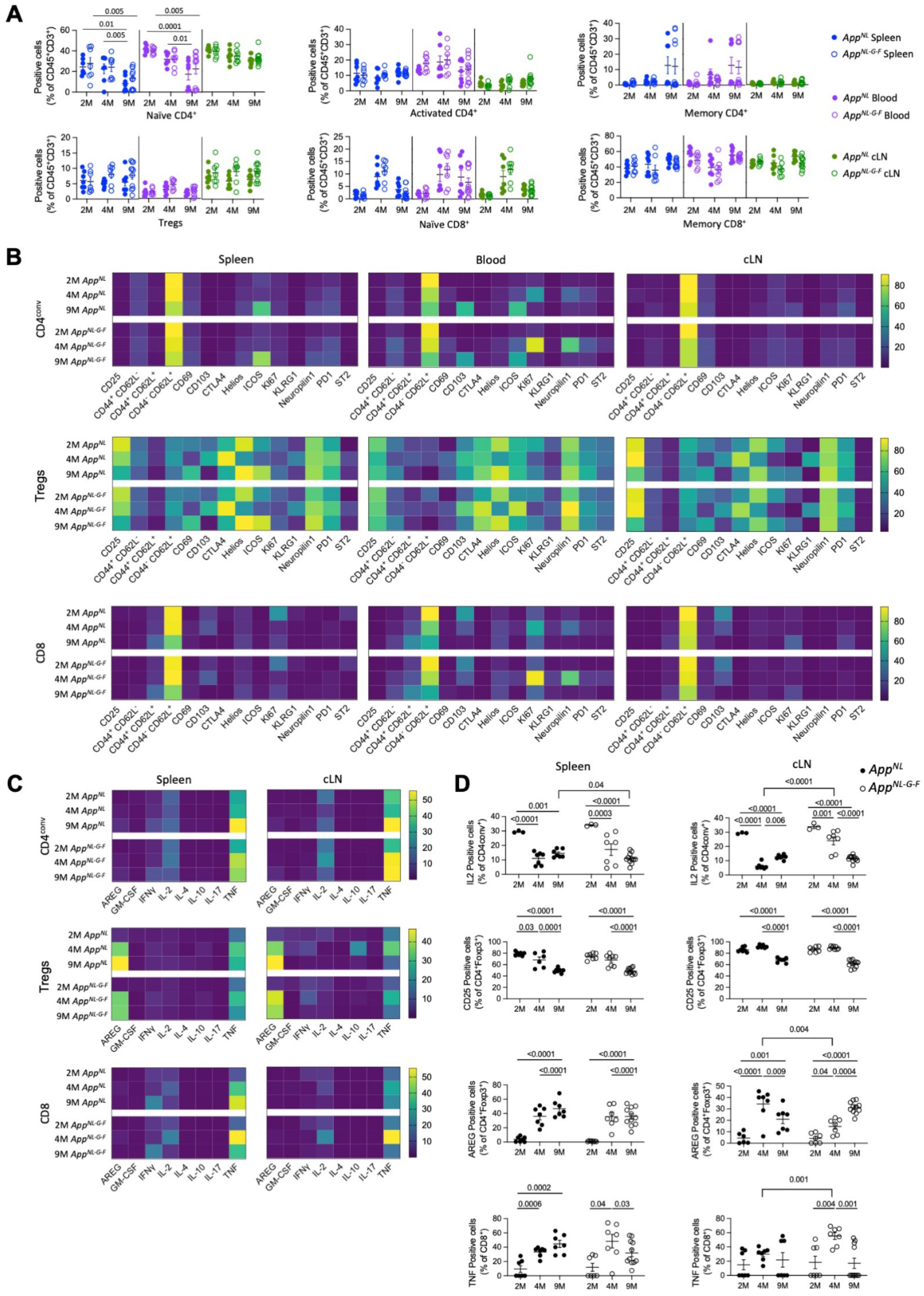
Cumulative age-dependent changes in the peripheral immunological compartments in *App*^*NL*^ and *App*^*NL-G-F*^ mice. *App*^*NL*^ and *App*^*NL-G-F*^ mice were assessed by high-dimensional flow cytometry for T cell immune profiles, at 2, 4 and 9 months of age. **A)** Frequency of main T cell populations in the spleen, blood, and cLN from *App*^*NL*^ and *App*^*NL-G-F*^ mice (2 months: n=7,7; 4 months: n=7,7; 9 months: n=7, 11), as defined by tSNE projection (Supplementary Figure 1). **B)** Heatmaps displaying activation marker expression in CD4^conv^, Tregs, and CD8 subsets in spleen, blood, and cLN, for each genotype and age. **C)** Heatmaps displaying expression of cytokines in CD4^conv^, Tregs, and CD8 subsets in spleen and cLN, for each genotype and age. **D**) Frequency of IL2^+^ cells within CD4^conv^ cells (2 months: n=3,3; 4 months: n=7,7; 9 months: n=7, 11), CD25^+^ cells within Tregs (2 months: n=7,7; 4 months: n=7,7; 9 months: n=7, 11), and AREG^+^ cells within Tregs (2 months: n=7,7; 4 months: n=7,7; 9 months: n=7, 11), measured by flow cytometry. Left column, Spleen; right column, cLN.

We next assessed the changes to T cell populations in the brain tissue. Major T cell subsets (**Figure 2A**) were quantified, demonstrating an age-dependent displacement of activated CD4 T cells and Tregs by activated CD8 T cells (**Figure 2B**). These effects were age-dependent rather than genotype-dependent, with few changes observed between *App*^*NL*^ and *App*^*NL-G-F*^ mice (**Supplementary Figure 5**). At the level of activation marker expression (**Figure 2C**), the only consistent change was in the age-dependent increase of CD44 expression on CD8 T cells (**Figure 2D**), reflecting the increase in the activated CD8 T cell population. Together, these results do not support a model of *App*^*NL-G-F*^ expression actively driving major immunological changes in the T cell compartment. However, the age-dependent changes are consistent with an alternative model, whereby the shifts in the T cell population in the older mouse (9 months) are permissive for a neuroinflammatory reaction precipitated by *App*^*NL-G-F*^ expression and subsequent Aβ accumulation. Under this latter model, *App*^*NL-G-F*^ expression in the young brain may be quiescent due to a protective immunological context, however normal “healthy” changes in phenotypes such as the IL2-Treg access (lower IL2 production by conventional CD4 T cells and lower CD25 expression on Tregs in the draining lymph nodes, lower Treg frequency in the brain tissue) may provide the immunological environment that allows *App*^*NL-G-F*^ expression to become pathogenic.

**Figure 2.**
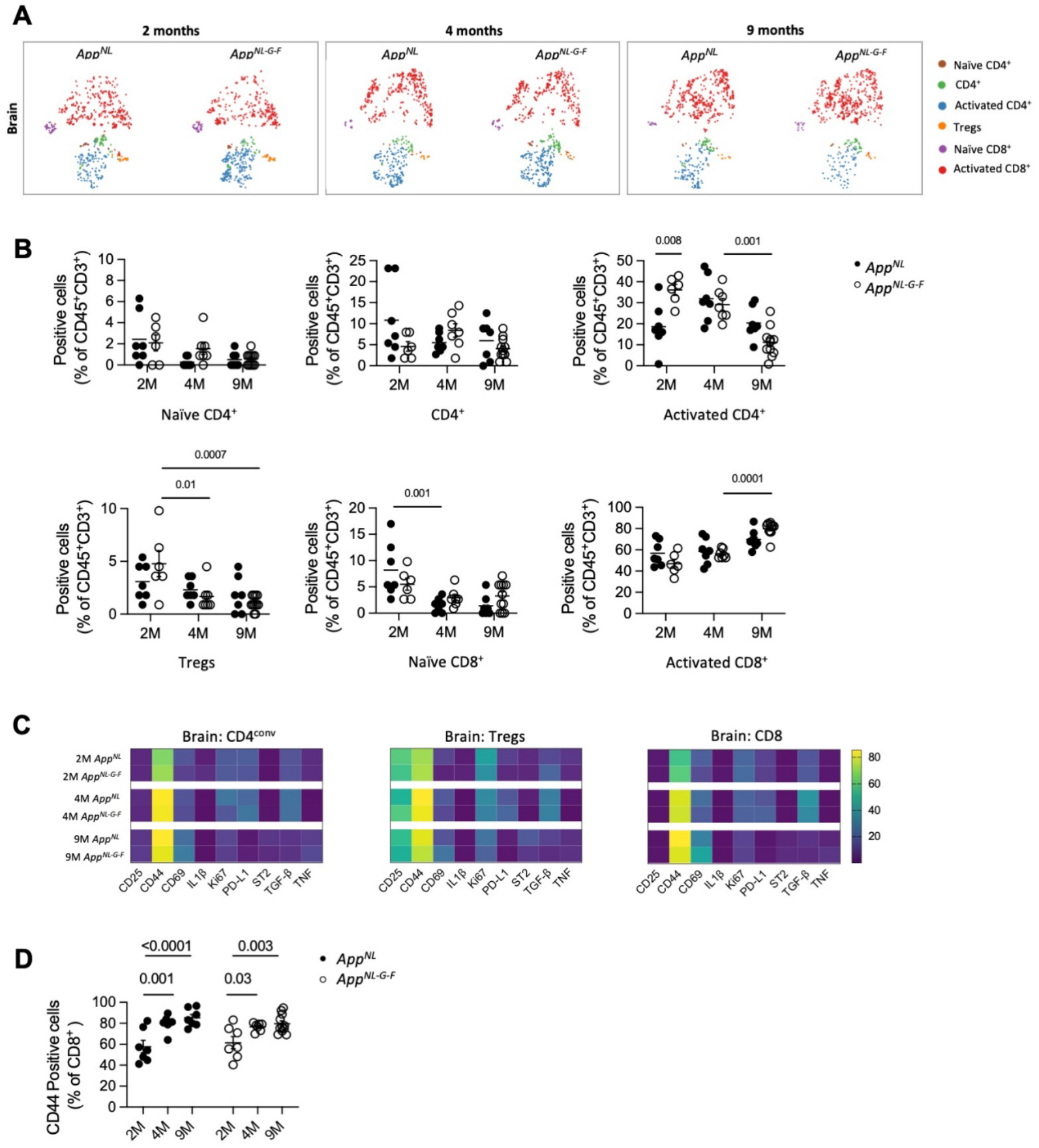
Cumulative age-dependent changes in the brain-resident immunological compartments in *App*^*NL*^ and *App*^*NL-G-F*^ mice. **A)** Perfused mouse brains from *App*^*NL*^ and *App*^*NL-G-F*^ mice were compared by high-dimensional flow cytometry to obtain the immune profiles of resident T cells, at 2 (n=7,7), 4 (n=7,7) and 9 (n=7,11) months of age. tSNE projection (4032 cells plotted) of the main T cell population clusters from concatenated samples. Immune populations were annotated based on expression of key protein markers, with **B)** cluster quantification. **C)** Heatmaps displaying the degree of expression for activation markers within the CD4^conv^, Tregs, and CD8 populations in the brains of *App*^*NL*^ and *App*^*NL-G-F*^ mice, at each age point. **D)** Frequency of CD44^+^ cells within the CD8 T cell population in the brain.

### Peripherally-biased expansion of Treg numbers via IL2 self-sufficiency does not alter pathology of AD

Our observation that the major identified age-dependent change observed in *App*^*NL*^ and *App*^*NL-G-F*^ mice is in the IL2-Treg axis is consistent with prior reports that IL2 or Treg treatment can prevent aspects of neurodegenerative phenotypes in AD model mice [8]. A limitation of prior studies is that treatment with either IL2 or Treg creates global changes, which do not allow a distinction between the role of Tregs in peripheral organs (e.g., regulation of priming events in the spleen or draining lymph nodes) versus the role of Tregs in the brain itself (e.g. promoting repair or creating an anti-inflammatory environment) to be determined. To separate these effects, we used a Cre-inducible IL2 transgene, allowing low levels of IL2 production to be initiated in a cell-dependent manner [23]. First, we used a *Foxp3*-Cre transgene to drive IL2 production in Tregs (Foxp3^Cre^RosaIL2 mice). By employing such a system, we effectively circumvent the normal dependence of Treg expansion on exogenous IL2 provision [23]. By crossing Foxp3^Cre^RosaIL2 mice to either *App*^*NL*^ and *App*^*NL-G-F*^ mice (**Figure 3A**), we generated a Treg-enriched environment in both lines (**Figure 3C)**. Due to the low levels of IL2 produced in this system, the non-Treg compartment remains unchanged, in both the periphery (spleen, blood, cLN) and brain (**Figure 3B**). The activation profile of both conventional CD4 and CD8 T cells remained unchanged by IL2 expression, in both periphery (**Supplementary Figures 6-7**) and brain (**Supplementary Figure 8**). By contrast, the Treg population increased in an age-dependent manner, with peripheral Tregs increasing by roughly 2-fold, 5-fold and 10-fold at 2, 4 and 9 months of age, respectively (**Figure 3C**). The expansion in the numbers of circulating Tregs occurs at the cost of the tissue-resident population [16]. This results in a substantial delay in Treg expansion in the brain tissue, with increases not observed until the mice are 9 months old (**Figure 3D**). This expansion process did not, however, substantially alter the activation profile of Tregs in either the brain (**Supplementary Figure 8**) or periphery (**Supplementary Figure 9**) at the 2, 4, and 9 month time-points. The system thereby provided *App*^*NL-G-F*^ -IL2 mice where dominant Treg expansion occurred in peripheral tissues, but not exclusively. Despite the substantial and early increase in peripheral Treg numbers in *App*^*NL-G-F*^ -IL2 mice, no change was observed in guanidine extracted Aβ (40, 42) measured using ELISA (**Figure 3E**), or in plaque accumulation measured by immunofluorescence (**Figure 3F**). Measurement of spatial learning in the Morris Water Maze showed clear cognitive decline in 9 month-old *App*^*NL-G-F*^ mice, with reduced preference for the target quadrant after learning (**Figure 3G**). *App*^*NL-G-F*^-IL2 mice showed only a minor effect no improvement in spatial cognitive learning over *App*^*NL-G-F*^ mice (**Figure 3G**), demonstrating that elevated peripheral Treg numbers were unable to prevent the cognitive decline observed in the *App*^*NL-G-F*^ model.

**Figure 3.**
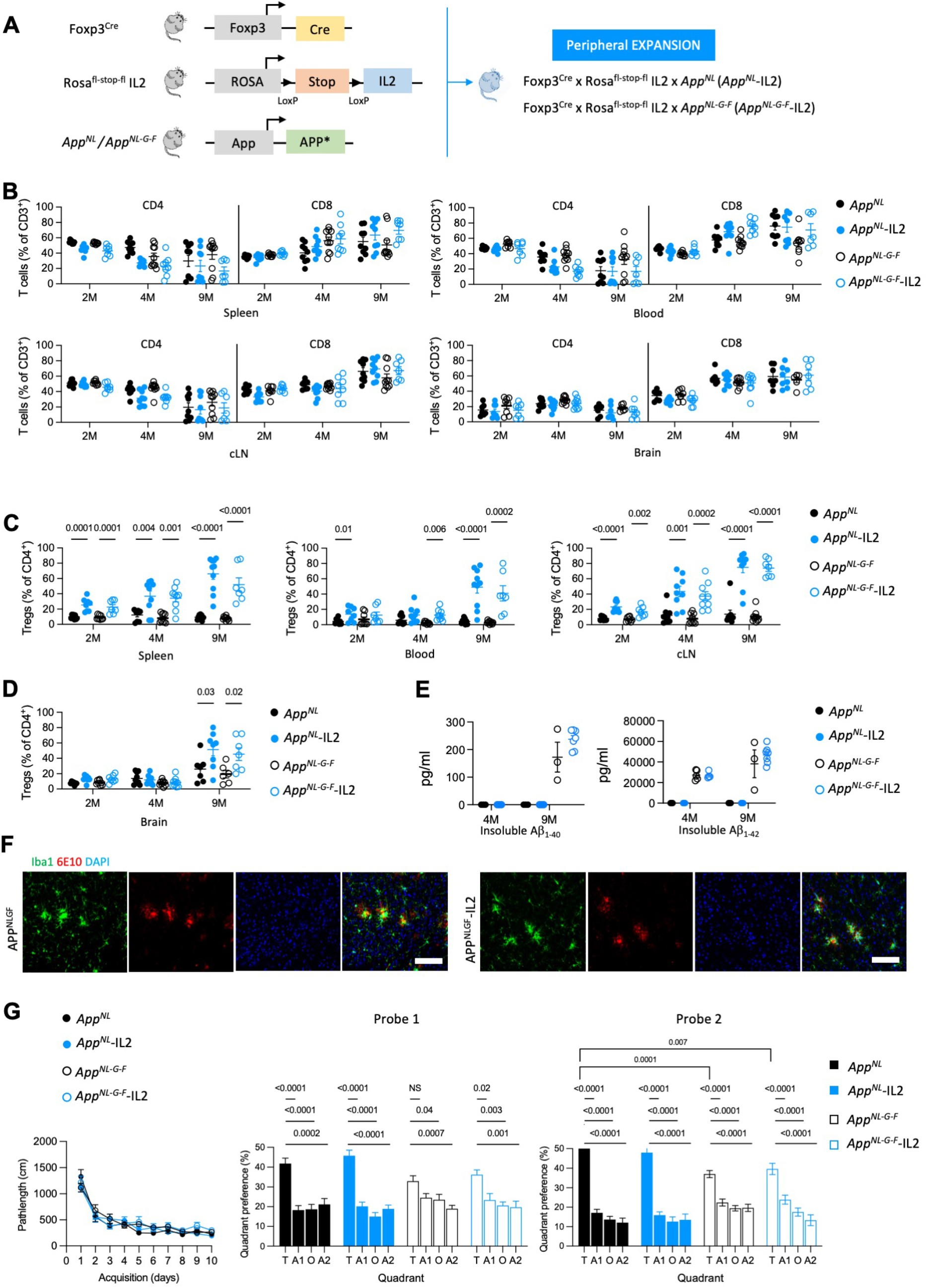
Peripherally-biased expansion of Treg numbers does not alter pathology of *App*^*NL-G-F*^ mice. **A)** Schematic of the transgenic system employed for IL2 overexpression in Tregs in *App*^*NL*^ and *App*^*NL-G-F*^ knock-in mice. **B)** Perfused mice (*App*^*NL*^, *App*^*NL*^ –IL2, *App*^*NL-G-F*^ and *App*^*NL-G-F*^ –IL2 lines) were compared by high-dimensional flow cytometry to determine the immune profiles in the T cell population. Frequency of CD4^conv^ and CD8 T cells in spleen, blood and cLN at 2 (n=7,8,8,7), 4 (n=9,9,9,9) and 9 (n=9,10,10,7) months of age. Frequency of CD4^conv^ and CD8 T cells in brain at 2 (n=7,8,8,7), 4 (n=9,9,9,9) and 9 (n=7,8,6,7) months of age. **C)** Frequency of Tregs in spleen, blood and cLN at 2 (n=7,8,8,7), 4 (n=9,9,9,9) and 9 (n=9,10,10,7) months of age. **D)** Frequency of Tregs in the brain at 2 (n=7,8,8,7), 4 (n=9,9,9,9) and 9 (n=7,8,6,7) months of age. **E)** ELISA measurements on levels of insoluble Aß1-40 (left) and Aß1-42 (right) extracted from whole cortex at 4 (n=5,5,5,4) and 9 (n=5,5,3,7) months of age. **F)** Representative cortical brain sections from 9 months old *App*^*NL-G-F*^ and *App*^*NL-G-F*^ -IL2 mice stained with anti-Iba1 and anti-Aß antibodies (6E10). Scale bar = 100 µm **G)** Spatial learning in the Morris Water Maze for *App*^*NL*^, *App*^*NL*^ –IL2, *App*^*NL-G-F*^ and *App*^*NL-G-F*^ –IL2 lines (n= 12,12,22,14). Mice were tested at 9 months of age. Left: Path length to finding the hidden platform; Middle: Probe tests after 5 days (probe 1); Right: after 10 days (probe 2) of acquisition.

### Brain-specific expansion of Treg numbers via brain-delivery of IL2 does not alter pathology of AD

Finally, we performed the reciprocal test, analyzing the effect of specific expansion of brain Tregs on APP-mediated pathology. Here we used the CamKII^Cre^ allele to trigger activation of the IL2 transgene specifically in CamKII^+^ neurons, previously demonstrated to drive a brain-specific expansion of Tregs [16]. By crossing CamKII^Cre^RosaIL2 mice to *App*^*NL-G-F*^ mice, we generated a brain-specific IL2 expression system in AD model mice (**Figure 4A**). Comparing *App*^*NL-G-F*^-CamKII^L2^ mice to *App*^*NL-G-F*^ mice, no changes were apparent in the frequency of conventional T cell populations, in either the peripheral tissues (blood, spleen, cLN) or the brain (**Figure 4B**). Likewise, activation profiles remained unchanged in both periphery (**Supplementary Figure 10-11**) and brain (**Supplementary Figure 12**). Treg frequency, by contrast, was normal in peripheral tissues, but was significantly increased (∼5-fold) in the brain, from 2 months of age (**Figure 4C**). Activation markers remained constant with IL2 treatment, demonstrating that the effect of IL2 was numerical expansion rather than alteration of phenotype (**Supplementary Figure 12-13**). This creates a context where we have a brain-specific increase in IL2 and Tregs from a young age, in the presence of a humanized, mutated *App* allele, designed to drive AD-like pathogenesis. Despite Tregs being elevated in the brain throughout the period of amyloid deposition, no significant changes were detected in the amount of insoluble Aβ1-40 and Aβ1-42 in the whole cortex (**Figure 4D**). Likewise, with regard to the development of behavioral abnormalities, *App*^*NL-G-F*^-CamKII^L2^ mice showed no improvement in the Morris Water Maze compared to *App*^*NL-G-F*^ mice **(Figure 4E**). Together with the results from the *App*^*NL-G-F*^-IL2 mice, this suggests that neither peripheral-biased nor brain-specific expansion of Tregs substantially alters pathology in the *App*^*NL-G-F*^ AD mouse model.

**Figure 4.**
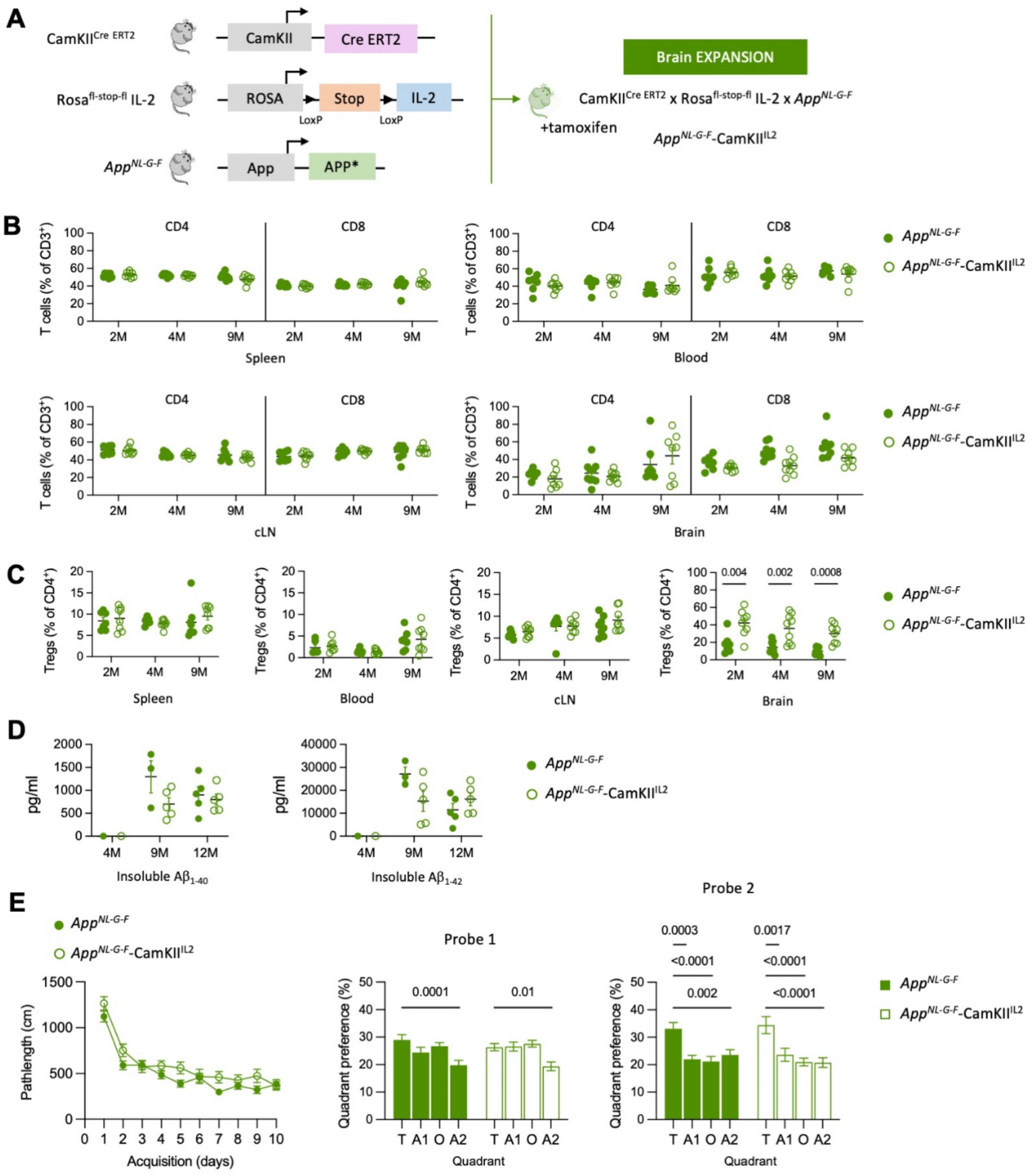
Brain-specific expansion of Treg numbers does not alter pathology in App knock-in mice. **A)** Schematic of the transgenic system employed for brain-specific IL2 expression in App knock-in mice. **B)** Perfused mice (*App*^*NL-G-F*^ and *App*^*NL-G-F*^ –CamK2^IL2^ lines) were compared by high-dimensional flow cytometry to determine the immune profiles in the T cell populations. Frequency of CD4^conv^ and CD8 T cells in spleen, blood and cLN at 2 (n=7,8), 4 (n=7,8) and 9 (n=8,8) months of age. Frequency of CD4^conv^ and CD8 T cells in brain at 2 (n=7,8), 4 (n=9,9) and 9 (n=8,8) months of age. **C)** Frequency of Tregs in spleen, blood and cLN at 2 (n=7,8), 4 (n=7,8) and 9 (n=8,8) months of age. Frequency of Tregs in the brain at 2 (n=7,8), 4 (n=9,9) and 9 (n=8,8) months of age. **D)** ELISA measurements on levels of insoluble Aß1-40 (left) and Aß1-42 (right) at 4 (n=4,4), 9 (n=3,5) and 12 (n=5,5) months of age. **E)** Spatial learning in the Morris Water Maze for *App*^*NL-G-F*^ and *App*^*NL-G-F*^ –CamK2^IL2^ lines (n= 26,20). Mice were tested at 9 months of age. Left: Path length to finding the hidden platform; Middle: Probe tests after 5 days (probe 1); Right: after 10 days (probe 2) of acquisition.

## Discussion

Unlike previous studies reporting either a protective effect of Tregs [8, 18] or a detrimental effect of Tregs [19, 20, 24] on disease progression in AD model mice, here we observed no effect of either peripheral-biased Treg expansion or brain-specific Treg expansion on cognitive decline and Amyloid beta aggregates. Of the potential explanations for the discrepancy in results, one is the use of different models of AD[25]. Previous studies on the role of Tregs on AD pathology have been performed in mouse models that overexpress the Aβ precursor protein (APP) in combination with different familial AD associated mutations, however multiple different models have been used. Alves *et al*, reporting a protective effect, used the APP/PS1 strain, which combine the Swedish APP mutation with the L166P PSEN1 mutation [8] and Baek *et al*, used the the 3xTg-AD mice containing three mutations associated with familial Alzheimer’s disease (APP Swedish, MAPT P301L, and PSEN1 M146V) [18]. Two studies have reported detrimental effects of Tregs, using the APP/PS1 mice [20], the 5XFAD mice [19], containing APP transgenes covering 5 AD-linked mutations and the L286V mutation in PSEN1,, Finally, the Dansokho study, which, like our study, reported no impact of Tregs on Aβ plaque load, used the APP/PS1 mice [21] and only a minor effect on behavioral tasks. Our study is the first to test the role of Tregs in the *App*^*NL -G-F*^ model. Unlike the other APP transgenic models previously used, *App*^*NL-G-F*^ is a single knock-in with expression driven by the endogenous promoter, and doers not harbor mutation in the PSEN1 gene, reducing artefacts caused by over-expression or off-target processing [22]. This is of particular importance in ascertaining the function of the immune system in AD processes, as human transgenic PSEN1 was found in splenocytes and off target processing could create a role for T cells in the model that does not represent the physiological process occurring in patients [26]. However, the presence of multiple mutations in the same APP gene, not observed in human patients, could in principle interact with each other in some cases that may not accurately represent clinical AD.

While the discrepancy between our result and previous studies observing an effect of Tregs may be due to the transgenic model used, it is notable that three separate studies using the APP/PS1 model demonstrated no effect, a beneficial effect and a detrimental effect of Tregs on plaque load [8, 20, 21]. These results are unlikely to be due to strain differences, and may reflect the diversity of approaches used to test the role of Tregs. Several studies have used depletion systems, either through PC61 anti-CD25 antibody or using the diphtheria toxin system in transgenic mice [19, 21]. These systems result in large-scale immune activation in addition to Treg depletion, and depending on the kinetics of treatment resurgent Treg number can even be locally elevated [19], confounding interpretation. Baek *et al* used the direct adoptive transfer of Tregs [19], which can transiently raise Treg numbers, although niche-sensing systems cause retraction of an overly abundant Treg population [17]. Finally, several studies have expanded endogenous Tregs, either through the provision of all-trans retinoic acid [19, 20] or IL2 [8, 21]. These approaches have the advantage of being therapeutically translatable, however there are limits to the degree to which mechanism of activity can be inferred. While all-trans retinoic acid expands Treg numbers [27], it has numerous effects on other aspects of the immune system [28] and nervous system, including amyloid processing[29]. Likewise, IL2 at low doses is a selective survival factor for Tregs, however at higher doses other cell types, including CD8 T cells, respond. Our study has the advantage that IL2 is being supplemented at doses low enough that only Tregs respond[16], with the cell lineage-mediated production of IL2 allowing segregation of peripheral and brain Treg functions.

At face value, the conclusions of our study do not support the use of IL2 or Treg therapy in AD. It is important, however, to consider that the lack of response may be driven by the limitations of the mouse model. *App*^*NL-G-F*^ mice only replicate a single aspect of AD disease progression, that driven by plaque formation, and can be considered to be models of preclinical AD, rather than of AD. *App*^*NL-G-F*^ mice are therefore a useful preclinical model to study processes such as glial responses to amyloid stress, but lack other important aspects of AD. The *App*^NL-G-F^ KI mice do, however, upregulate multiple neuroinflammation-related genes in common with AD patients genes (*C4a/C4b, Cd74, Ctss, Gfap, Nfe212, Phyhd1, S100b, Tf, Tgfbr2* and *Vim*) and AD risk factor genes (A*bi3, Apoe, Bin2, Cd33, Ctsc, Dock2, Fcer1g, Frmd6, Hck, Inpp5D, Ly86, Plcg2, Trem2* and *Tyrobp*)[22, 24, 30]. Importantly, the mice do not exhibit tau pathology or neurodegeneration [22], a key phenotype in patients [31], suggesting that can be useful as preclinical AD model to investigate the pathological role of amyloid-associated neuroinflammation The lack of tau pathology is particularly pertinent as studies of human AD postmortem brains indicates that T cell infiltration correlates with tau pathology rather than with amyloid plaques [32]. The APP^NLGF^ mouse model also progresses independent of the adaptive immune system, in Rag-deficient mice [33], contradicting genetic association of human AD with adaptive immunity [6, 7]. We therefore provide the more nuanced conclusion that the current results do not support the use of IL2 or Treg therapy in AD patients where the primary driver is APP, without discounting potential efficacy in AD patients where tau pathology is a primary driver. Indeed, we have previously demonstrated that brain-specific delivery of IL2 prevents cognitive decline in APP-independent mouse models of traumatic brain injury [16] and old age [34], validating the use of IL2 or Treg therapy in dementia more generally. We therefore echo the call that any potential clinical trials should take into account the heterogeneity of AD disease, with stratification based on genotype and tau pathology increasing the potential for efficacious responses.

## Acknowledgements

The work was supported by the Alzheimer’s Association AARG (to A.L.), the VIB, an ERC Consolidator Grant TissueTreg (to A.L.), an ERC Proof of Concept Grant TreatBrainDamage (to A.L.), FWO Research Grant 1503420N (to EP), an SAO-FRA pilot grant (20190032, to E.P.), an ERC Starting Grant AstroFunc (to M.G.H.), the ERC Proof of Concept Grant AD-VIP (to M.G.H.), FWO Research Grant 1513616N (to M.G.H.), Thierry Latran Foundation Grant SOD-VIP (to M.G.H.), SAO-FRA pilot grant (14006) (to M.G.H.), ERNAET Chair NCBio (to M.G.H.), and the Biotechnology and Biological Sciences Research Council through Institute Strategic Program Grant funding BBS/E/B/000C0427 and BBS/E/B/000C0428, and the Biotechnology and Biological Sciences Research Council Core Capability Grant to the Babraham Institute. E.P. was supported by a fellowship from the FWO. The authors acknowledge the important contributions of the KUL FACS Core.

