## Supplementary material for "The *App*^*NL-G-F*^ mouse model of Alzheimer’s disease is refractory to regulatory T cell treatment": Suppl material

### Methods

#### *Mice*

Human amyloid precursor protein (hAPP) knock-in (KI) mice containing the Swedish, Iberian, and Arctic mutations included (*App*$^{NL\text{-}G\text{-}F}$) were used on a C57BL/6 J background [1]. Foxp3-Cre transgenic mice [2] and αCamKII-CreERT2 transgenic mice [3] were used on the C57BL/6 background. RosaIL2 mice were generated through the insertion of a cassette containing a floxed-STOP sequence followed by an IL2-IRES-GFP sequence into the Rosa26 locus, using the endogenous Rosa26 promoter [4], and were used on the C57BL/6 background. Both male and female mice were used in this study. Mice were age-matched and tested at 2, 4, and 9 months of age. Tamoxifen (Sigma T5648) was solubilized in corn oil (Sigma) at 10 mg/ml. Five to seven weeks old αCamKII-CreERT2 mice were injected 3 times, via intraperitoneal injection, at 48 h intervals using a dose of 100 mg/kg body weight. Mice were housed under SPF conditions, under a 12-hour light/dark cycle in a temperature and humidity-controlled room with *ad libitum* access to food and water. All animal procedures were approved by the KU Leuven Animal Ethics Committee (P124/2019), and were consistent with European guidelines.

#### *Flow cytometry*

Mice were deeply anaesthetized with intraperitoneal injection of a ketamine (87 mg/kg), xylazine (13 mg/kg) mixture. Blood was collected from the right ventricle prior to transcardial perfusion with ice cold PBS, and further processed through red blood cell lysis. Single-cell suspensions from lymphoid organs were prepared by mechanical dissociation; single-cell suspensions from brain tissue were prepared by digestion for 30 minutes at 37ºC with 1 mg/ml collagenase IV (Thermo Fisher), 300 µg/ml hyaluronidase (Sigma-Aldrich) and 40 µg/ml DNase I (Sigma-Aldrich) in RPMI 1640 supplemented with 2 mM $MgCl_2$, 2mM $CaCl_2$, 20% FBS and 2 mM HEPES (Gibco), followed by mechanical disruption, filtration (through 100 µm mesh) and enrichment for leukocytes by gradient centrifugation (40% Percoll GE Healthcare, 600 x g, 10 min). Non-specific binding was blocked using 2.4G2 supernatant. To assess intracellular cytokine production, cells were cultured for 4h in the presence of phorbol mysristate acetate (1 µg/ml, Sigma-Aldrich), ionomycin (1 µg/ml, Sigma-Aldrich), and brefeldinA (BD). Cells were fixed and permeabilized with the eBioscience Foxp3 staining kit (eBioscience). Cellular phenotypes were assessed using high parameter flow cytometry panels, containing markers to identify cell types and markers to assess activation states. Data were acquired on a BD FACSymphony, with panels covering (i) CD45, CD4, CD8, CD3, CD19, NK1.1, Foxp3, eBioscience™ Fixable Viability Dye eFluor™ 780, CD103, CD62L, CD25, Neuropilin, ST2, PD-1, KLRG1, Helios, CD69, ICOS, CD44, and Ki67 or (ii) CCR6, CD80, TCRγδ, CD45, Foxp3, MHCII, eBioscience™ Fixable Viability Dye eFluor™ 780, IL1β, CD25, Ly6G, ST2, CX3CR1, PD-L1, TNF, CD44, Ki67, CD4, Ly6C, TrkB, CD19, CD69, CD8α, LAMP1, CD64, CD11b, CD3, or (iii) Foxp3, eBioscience™ Fixable Viability Dye eFluor™ 780, IL5, IL6, IL17, CD4, IFNγ, CD8α, TNFα, CD3, Amphiregulin, IL10, IL4, CD11b, CD19, GM-CSF, TCRγδ, pro-IL1β, TCRβ, IL2, NK1.1. For brain panels, cells obtain from the whole brain were analysed. Data was compensated using AutoSpill [5].

tSNE, FlowSOM and heatmap analysis were performed in RStudio (version 1.4.1717) using an in-house script [5]. FlowSOM clusters are formed based on multi-marker similarity in a non-supervised manner. Clusters were annotated based on post-clustering comparison of marker expression, aligning the unique marker profile of each cluster to literature-based nomenclature. Key annotations for T cell clusters included CD4 naïve ($CD3^+CD4^+CD62L^{hi}CD44^{low}$), CD4 activated ($CD3^+CD4^+CD62L^{low}CD44^{high}$), CD4 memory ($CD3^+CD4^+CD62L^{hi}CD44^{high}$), Tregs ($CD3^+CD4^+Foxp3^+$), CD8 naïve ($CD3^+CD8^+CD62L^{hi}CD44^{low}$), CD8 activated ($CD3^+CD8^+CD62L^{low}CD44^{high}$), and CD8 memory ($CD3^+CD8^+CD62L^{hi}CD44^{high}$).

*Immunostaining*

Mice were deeply anaesthetized using intraperitoneal injection of a ketamine (87 mg/kg) / xylazine (13 mg/kg) mixture and transcardially perfused with PBS followed by 4% buffered formalin solution. The brain was removed and fixed in 10% buffered formalin solution overnight and stored in 30% sucrose until preservation in tissue freezing medium (Shandon™ Cryomatrix™ embedding resin, Thermo Scientific), and stored at -80°C. Sections (50 µm) were washed 15 minutes in 50 mM $NH_4Cl$/PBS and pre-blocked with 10% normal donkey serum in 0.5% Triton-X-100/PBS for 1h at room temperature. Sections were incubated overnight at 4ºC with primary antibodies directed against Iba1 (1:1000, 014-19741, Wako) and 6E10 Abeta (1:1000, SIG-39320-1000, Covance Signet). Subsequently, the sections were incubated for 90 minutes at room temperature with appropriate fluorophore-conjugated secondary antibodies (Thermo Scientific). After each antibody incubation, slices were washed 3 times for 10 minutes with 0.1% Triton-X-100/PBS. All sections were incubated with DAPI (1:1000) for 15 min before final mounting on microslide slides using ProlongGold (Invitrogen). Images were obtained using a Nikon A1R Eclipse Ti confocal (Plan Apo 20X), or a Zeiss Axioscan Z.1 slide-scanner (20X Plan-Apochromat/NA 0.8) equipped with a Hamamatsu Orca Flash 4.0 V3 camera. Image processing was performed using ImageJ (https://imagej.nih.gov/ij/download.html).

*ELISA*

Soluble and insoluble fractions of Aβ were extracted from cortex and hippocampus, as previously described [6] 1368. The total protein content of the samples containing the soluble and insoluble Aβ fractions was determined using a modified Lowry-Peterson assay. The concentration of $A\beta_{1-40}$ or $A\beta_{1-42}$ in tissue samples was determined using standard sandwich ELISAs. Monoclonal antibodies JRFcAβ40/28 and JRFcAβ42/26, which recognize the C-terminal ends of Aβ species terminating at amino acids 40 or 42 respectively, were used for capture. HRP-conjugated JRFAβN/25 antibody, which recognizes the first seven N-terminal amino acids of human Aβ, was used as a detection antibody. Synthetic human $A\beta_{1-40}$ and $A\beta_{1-42}$ peptides were used to generate standard curves. Antibodies were kindly provided by Johnson and Johnson. Absorbance was measured at 450 nm in a Perkin Elmer EnVision 2103 Multilabel reader.

*Morris Water Maze*

Morris Water Maze behavioral experiments were performed in 9 months old mice. Mice were habituated to their new environment for at least 7 days and tests were conducted during the light phase of their activity cycle. Tests were performed and analyzed by an observer blind to the experimental group. Spatial learning and cognitive flexibility were tested in the hidden platform Morris Water Maze. A circular pool (150 cm diameter) was filled with opacified (0.01% Acusol OP301, Dow Chemicals) water (26 ± 1°C). The platform (15 cm diameter) was hidden 1 cm underneath the surface of the water. For spatial learning, the mice were trained for 10 days to a fixed platform position [7, 8]. To evaluate reference memory, probe trials (100 seconds) were conducted on days 6 and 11 during acquisition learning. During probe trials, floater mice were excluded. The escape platform was removed from the pool and mice were allowed to explore the maze for 100 seconds. Swim paths were tracked with Ethovision software (Noldus).

*Statistics*

Comparisons between two groups were performed using unpaired two-tailed Student's t tests. Post hoc Holm's or Sidak's multiple comparisons tests were performed, when required. Two-way ANOVA was used, when appropriate. The value of n reported within figure legends represents the number of animals. Values are represented as mean ± SEM, with differences considered significant when $p < 0.05$.

**Supplementary Figure 1. Flow cytometry analysis of peripheral T cells in APP knock-in mice.** Perfused mouse brains from $App^{NL}$ and $App^{NL\text{-}G\text{-}F}$ mice were compared by high-dimensional flow cytometry for T cell immune profiles, at 2 (n=7,7), 4, (n=7,7) and 9 (n=7, 11) months of age. tSNE projection (27,638 cells plotted) of the main T cell population clusters. Immune populations were annotated based on key markers.

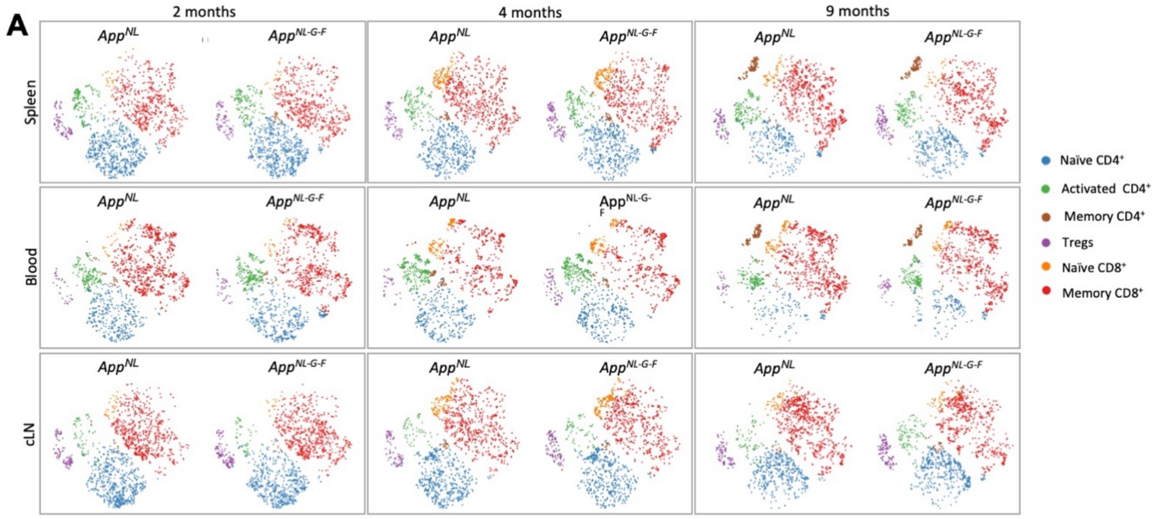

**Supplementary Figure 2. Flow cytometry analysis of peripheral Tregs in APP knock-in mice.** Perfused mice from $App^{NL}$ and $App^{NL\text{-}G\text{-}F}$ mice were compared by high-dimensional flow cytometry to ascertain the immune profile of T cells. Frequency of CD25, CD44$^+$CD62L$^-$, CD44$^+$CD62L$^+$, CD44$^-$CD62L$^+$, CD69, CD103, CTLA4, Helios, ICOS, Ki67, KLRG1, Neuropilin1, PD1 and ST2 expression within the peripheral Treg population in spleen at **A)** 2 (n=7,7), **B)** 4 (n=7,7) and **C)** 9 (n=7,11) months of age; in cLN at **D)** 2 (n=7,7), **E)** 4 (n=7,7) and **F)** 9 (n=7,11) months of age; and in the blood at **G)** 2 (n=7,7), **H)** 4 (n=7,7) and **I)** 9 (n=7,11) months of age.

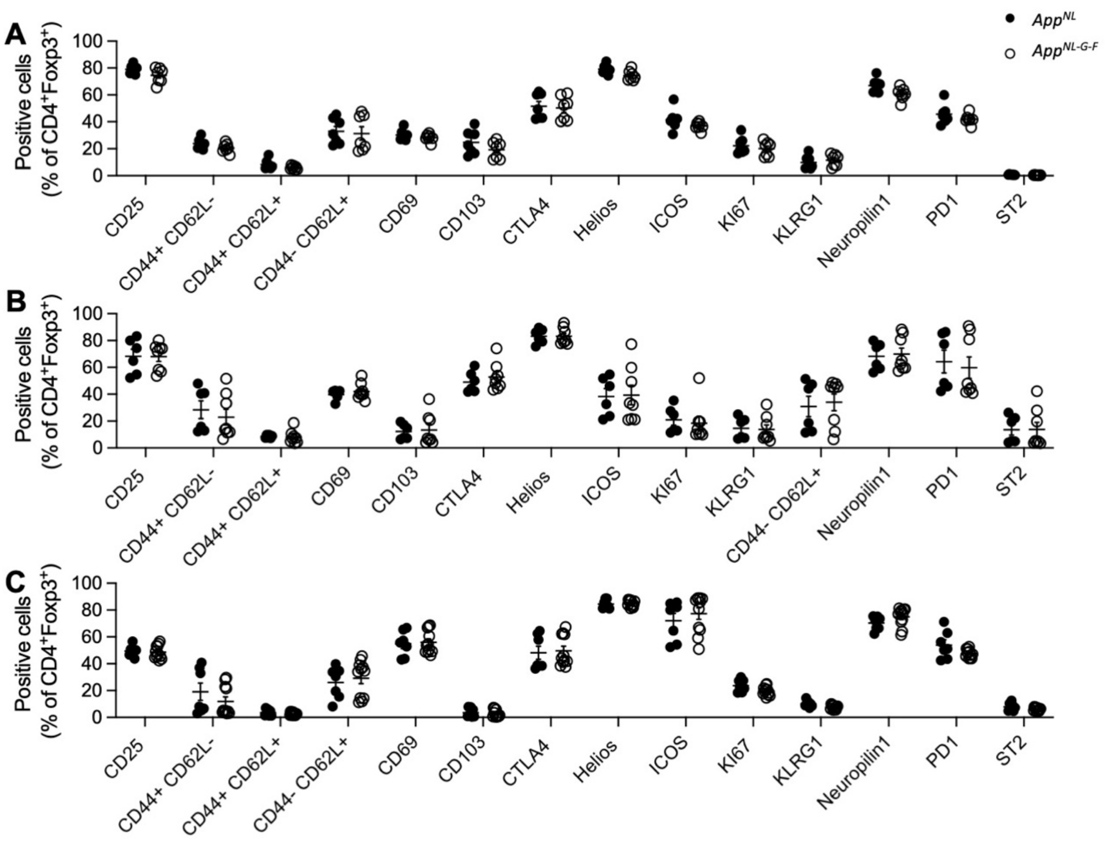

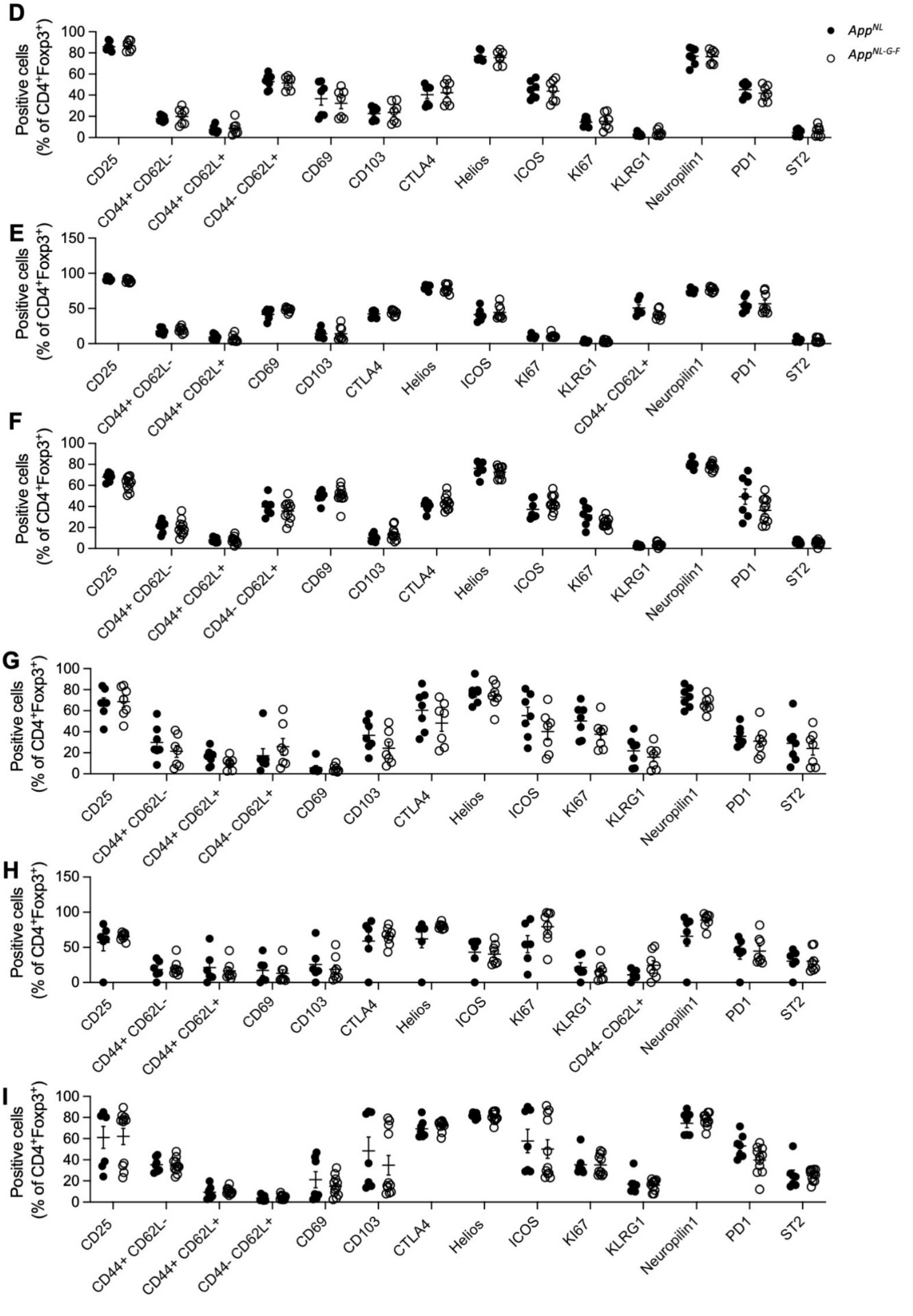
D
Positive cells (% of CD4+Foxp3+)
App^NL
App^NL-G-F
CD25
CD44+ CD62L-
CD44+ CD62L+
CD44- CD62L+
CD69
CD103
CTLA4
Helios
ICOS
KI67
KLRG1
Neuropilin1
PD1
ST2
E
Positive cells (% of CD4+Foxp3+)
CD25
CD44+ CD62L-
CD44+ CD62L+
CD69
CD103
CTLA4
Helios
ICOS
KI67
KLRG1
CD44- CD62L+
Neuropilin1
PD1
ST2
F
Positive cells (% of CD4+Foxp3+)
CD25
CD44+ CD62L-
CD44+ CD62L+
CD44- CD62L+
CD69
CD103
CTLA4
Helios
ICOS
KI67
KLRG1
Neuropilin1
PD1
ST2
G
Positive cells (% of CD4+Foxp3+)
CD25
CD44+ CD62L-
CD44+ CD62L+
CD44- CD62L+
CD69
CD103
CTLA4
Helios
ICOS
KI67
KLRG1
Neuropilin1
PD1
ST2
H
Positive cells (% of CD4+Foxp3+)
CD25
CD44+ CD62L-
CD44+ CD62L+
CD69
CD103
CTLA4
Helios
ICOS
KI67
KLRG1
CD44- CD62L+
Neuropilin1
PD1
ST2
I
Positive cells (% of CD4+Foxp3+)
CD25
CD44+ CD62L-
CD44+ CD62L+
CD44- CD62L+
CD69
CD103
CTLA4
Helios
ICOS
KI67
KLRG1
Neuropilin1
PD1
ST2

**Supplementary Figure 3. Flow cytometry analysis of peripheral $CD4^{conv}$ T cells in APP knock-in mice.** Perfused mice from $App^{NL}$ and $App^{NL\text{-}G\text{-}F}$ mice were compared by high-dimensional flow cytometry to ascertain the immune profile of T cells (?). Frequency of CD25, $CD44^+CD62L^-$, $CD44^+CD62L^+$, $CD44^-CD62L^+$, CD69, CD103, CTLA4, Helios, ICOS, Ki67, KLRG1, Neuropilin1, PD1 and ST2 expression within the peripheral $CD4^{conv}$ T cell population in spleen at **A)** 2 (n=7,7), **B)** 4 (n=7,7) and **C)** 9 (n=7,11) months of age; in cLN at **D)** 2 (n=7,7), **E)** 4 (n=7,7) and **F)** 9 (n=7,11) months of age; and in the blood at **G)** 2 (n=7,7), **H)** 4 (n=7,7) and **I)** 9 (n=7,11) months of age.

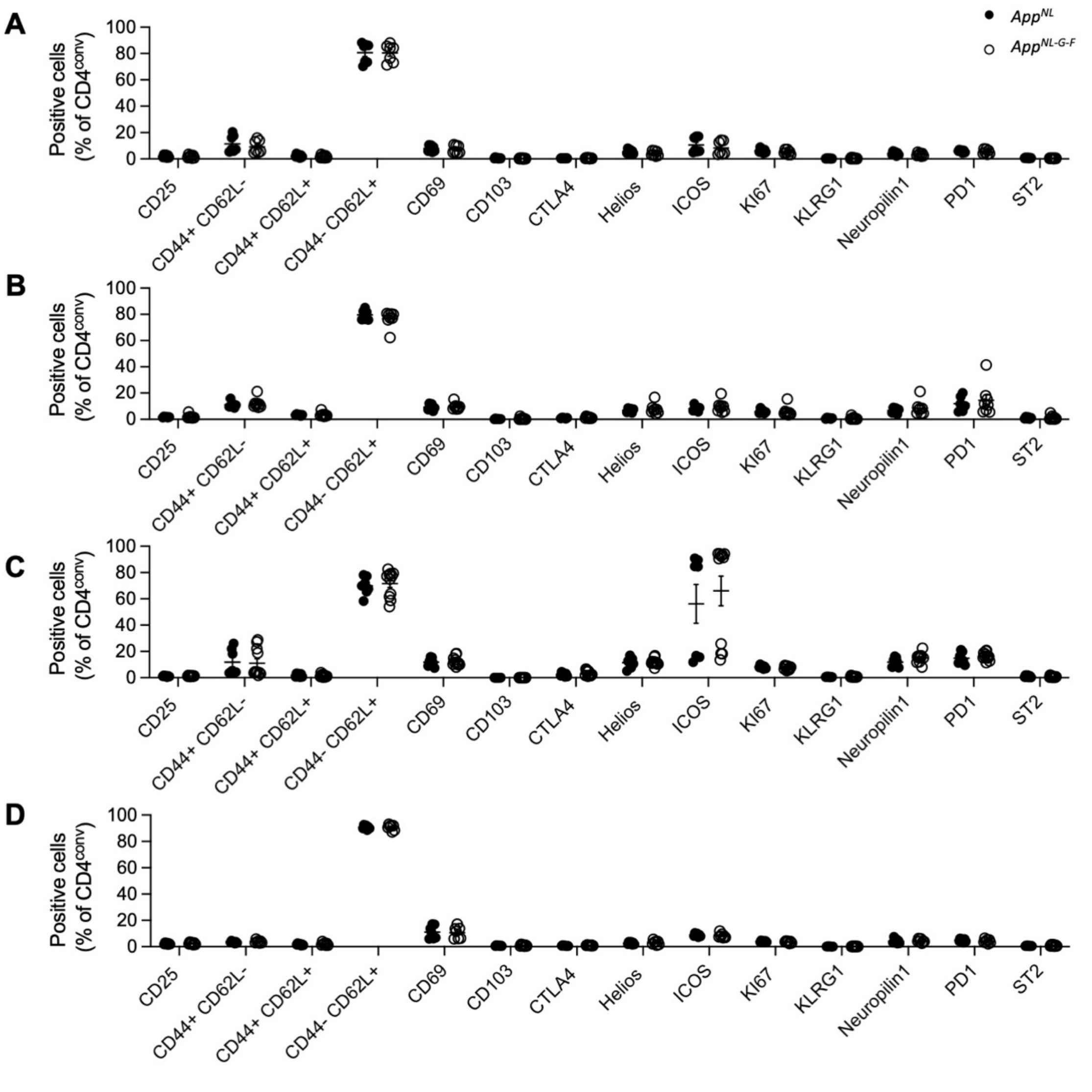

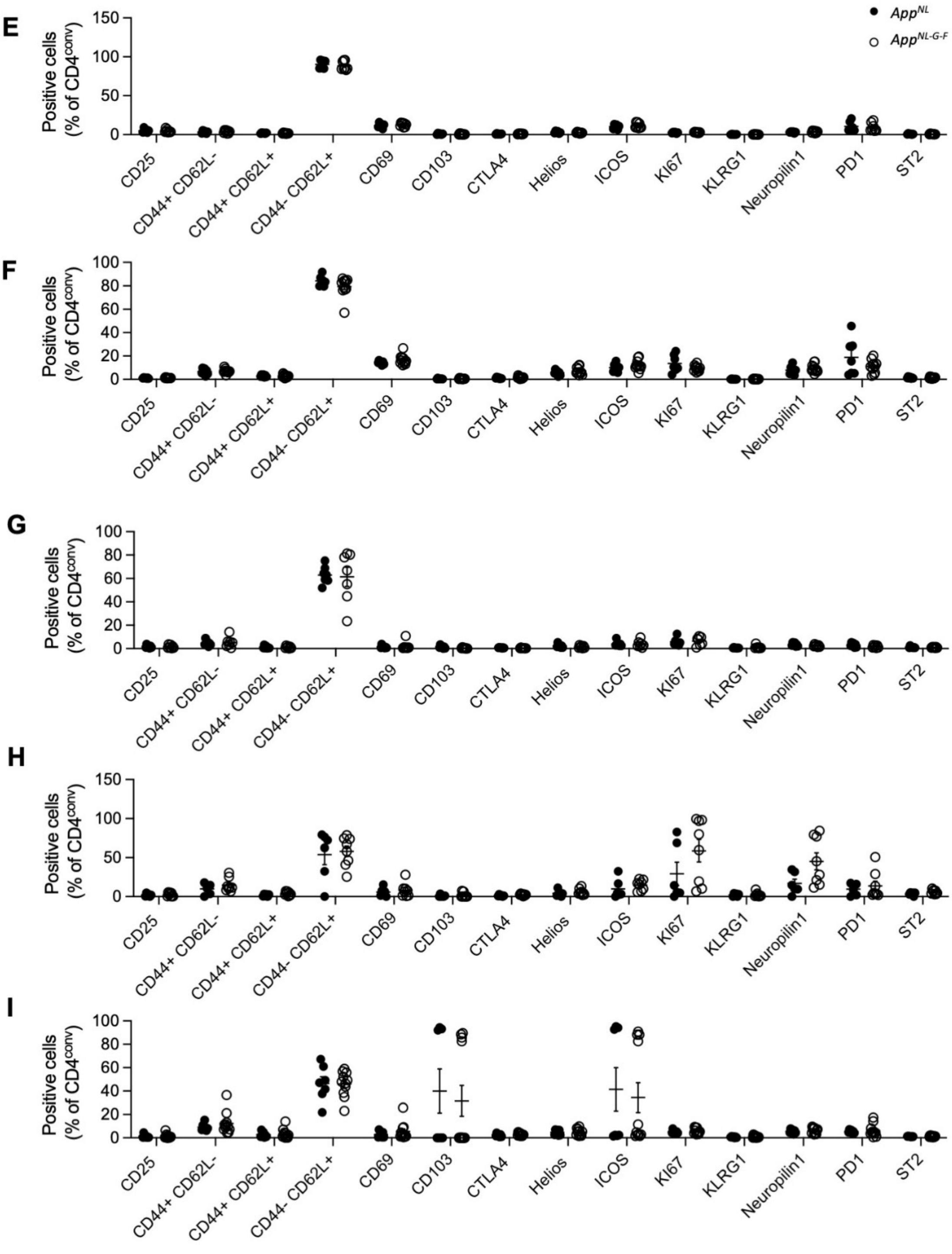
E
F
G
H
I
Positive cells
(% of CD4conv)
AppNL
AppNL-G-F
CD25
CD44+ CD62L-
CD44+ CD62L+
CD44- CD62L+
CD69
CD103
CTLA4
Helios
ICOS
KI67
KLRG1
Neuropilin1
PD1
ST2

**Supplementary Figure 4. Flow cytometry analysis of peripheral CD8 T cells in APP knock-in mice.** Perfused mice from *App*$^{NL}$ and *App*$^{NL\text{-}G\text{-}F}$ mice were compared by high-dimensional flow cytometry to ascertain the immune profile of T cells. Frequency of CD25, CD44$^{+}$CD62L$^{-}$, CD44$^{+}$CD62L$^{+}$, CD44$^{-}$CD62L$^{+}$, CD69, CD103, CTLA4, Helios, ICOS, Ki67, KLRG1, Neuropilin1, PD1 and ST2 expression within the peripheral CD8 T cell population in spleen at **A)** 2 (n=7,7), **B)** 4 (n=7,7) and **C)** 9 (n=7,11) months of age; in cLN at **D)** 2 (n=7,7), **E)** 4 (n=7,7) and **F)** 9 (n=7,11) months of age; and in the blood at **G)** 2 (n=7,7), **H)** 4 (n=7,7) and **I)** 9 (n=7,11) months of age.

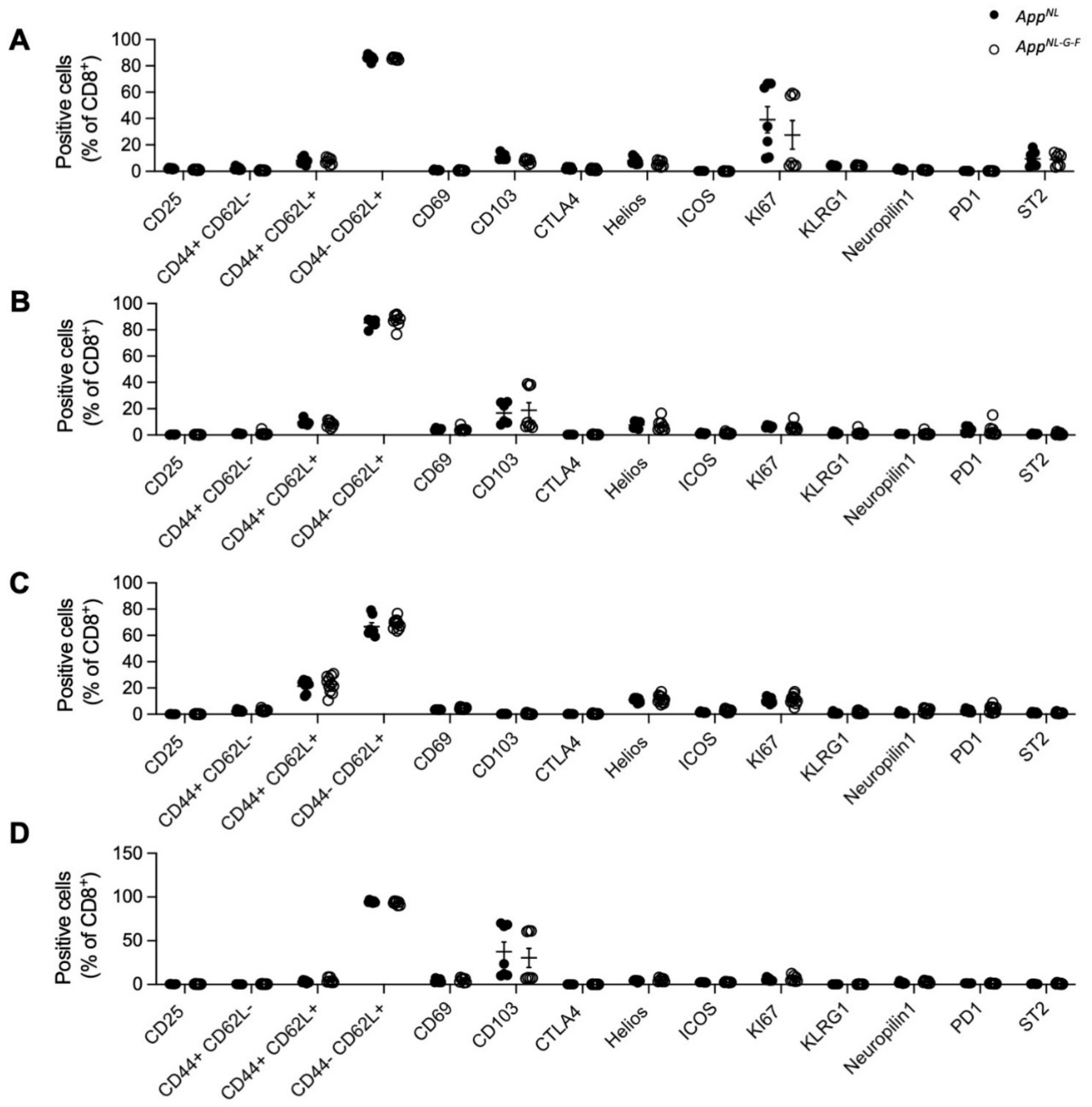

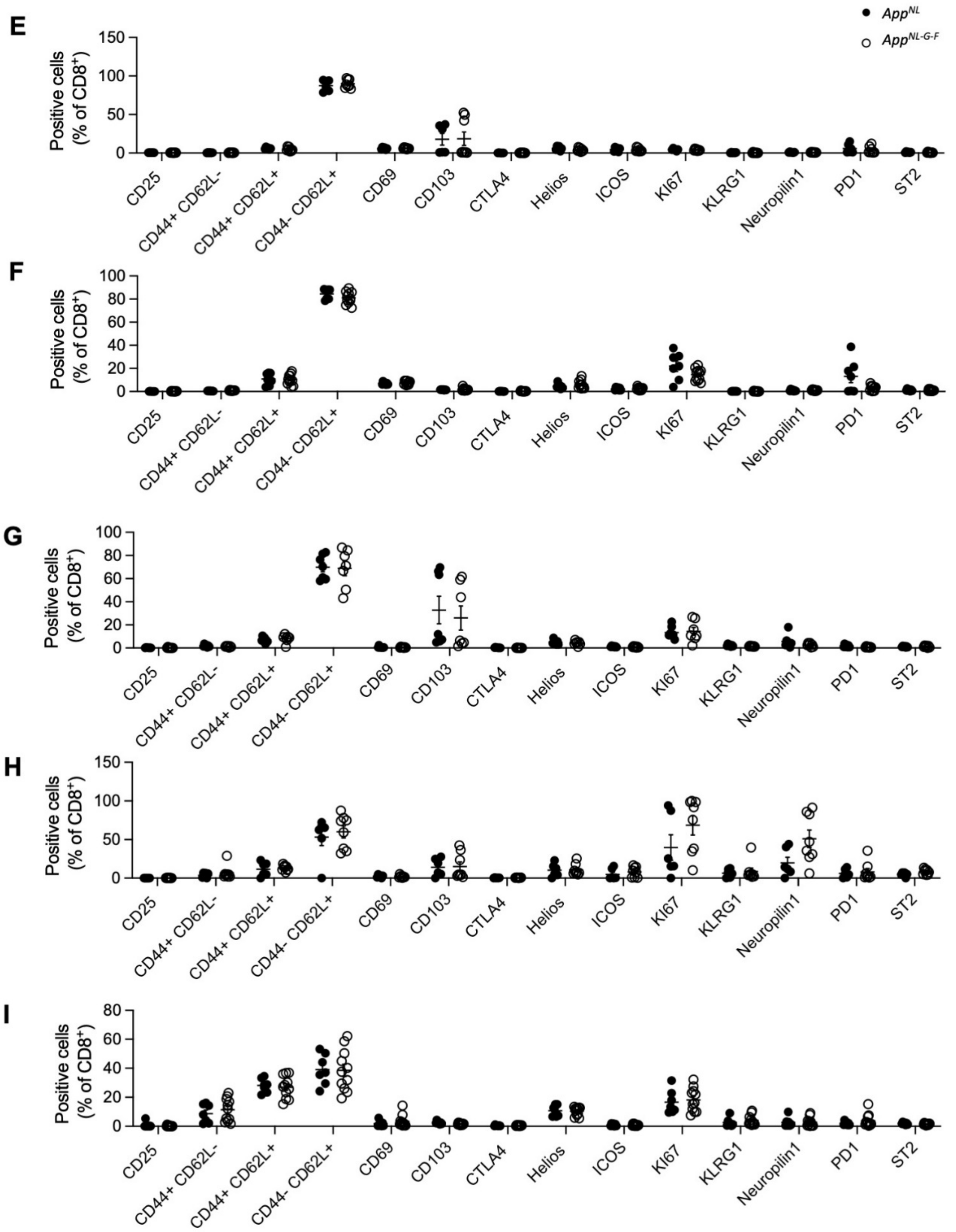
E
F
G
H
I
App^NL
App^NL-G-F
Positive cells (% of CD8+)
150
100
80
60
50
40
20
0
CD25
CD44+ CD62L-
CD44+ CD62L+
CD44- CD62L+
CD69
CD103
CTLA4
Helios
ICOS
KI67
KLRG1
Neuropilin1
PD1
ST2

**Supplementary Figure 5. Flow cytometry analysis of brain T cells in APP knock-in mice.** Perfused brains from *App*$^{NL}$ and *App*$^{NL\text{-}G\text{-}F}$ mice were compared by high-dimensional flow cytometry to determine the immune profile of brain T cells. Frequency of CD25, CD44, CD69, IL-1ß, Ki67, PD-L1, ST2, TGF-ß, and TNF expression within the brain Tregs population at **A)** 2 (n=7,7), **B)** 4 (n=7,7) and **C)** 9 (n=7,11) months of age; CD4$^{conv}$ T cells at **D)** 2 (n=7,7), **E)** 4 (n=7,7) and **F)** 9 (n=7,11) months of age; and within brain CD8 T cells at **G)** 2 (n=7,7), **H)** 4 (n=7,7) and **I)** 9 (n=7,11) months of age.

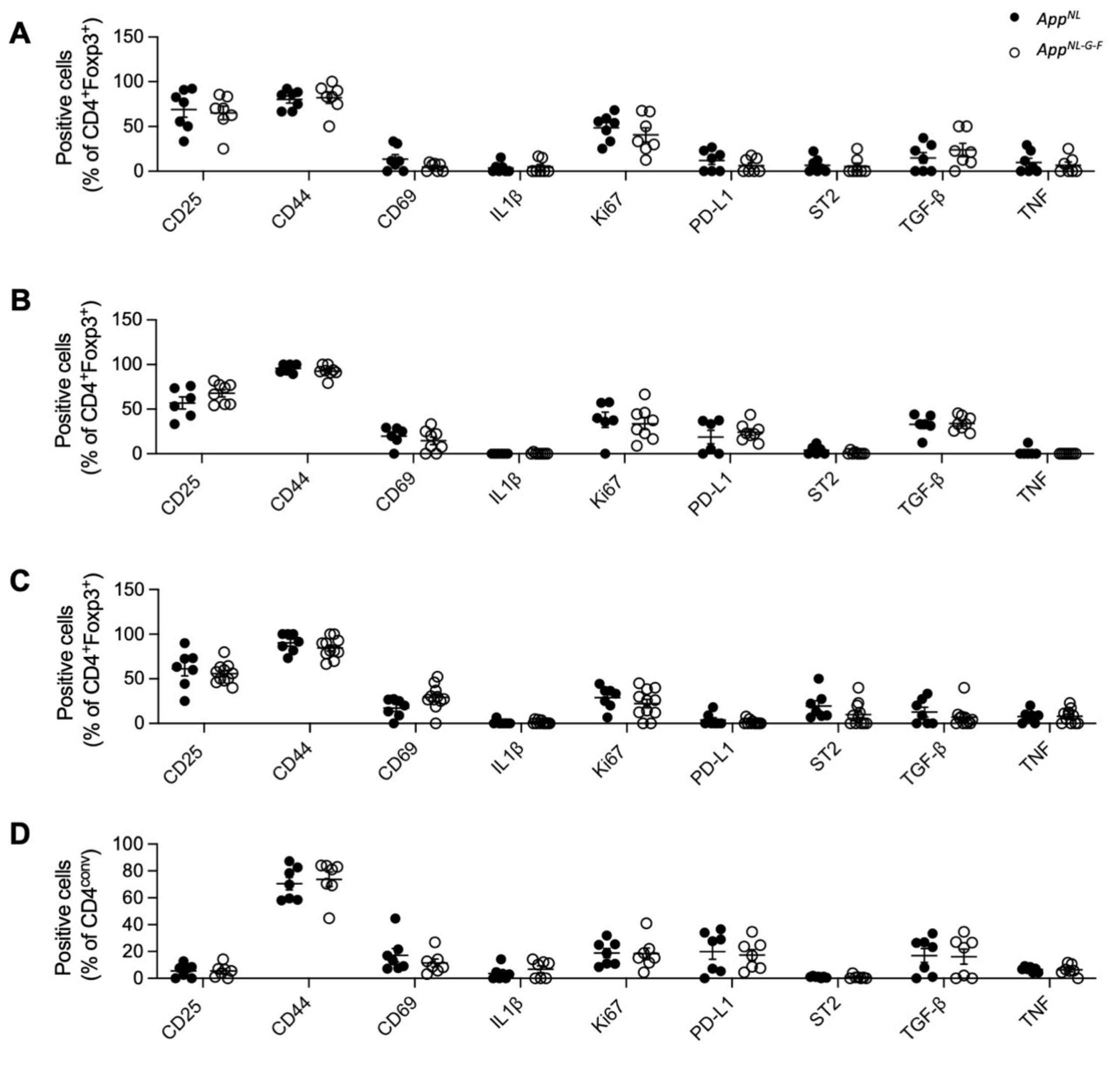

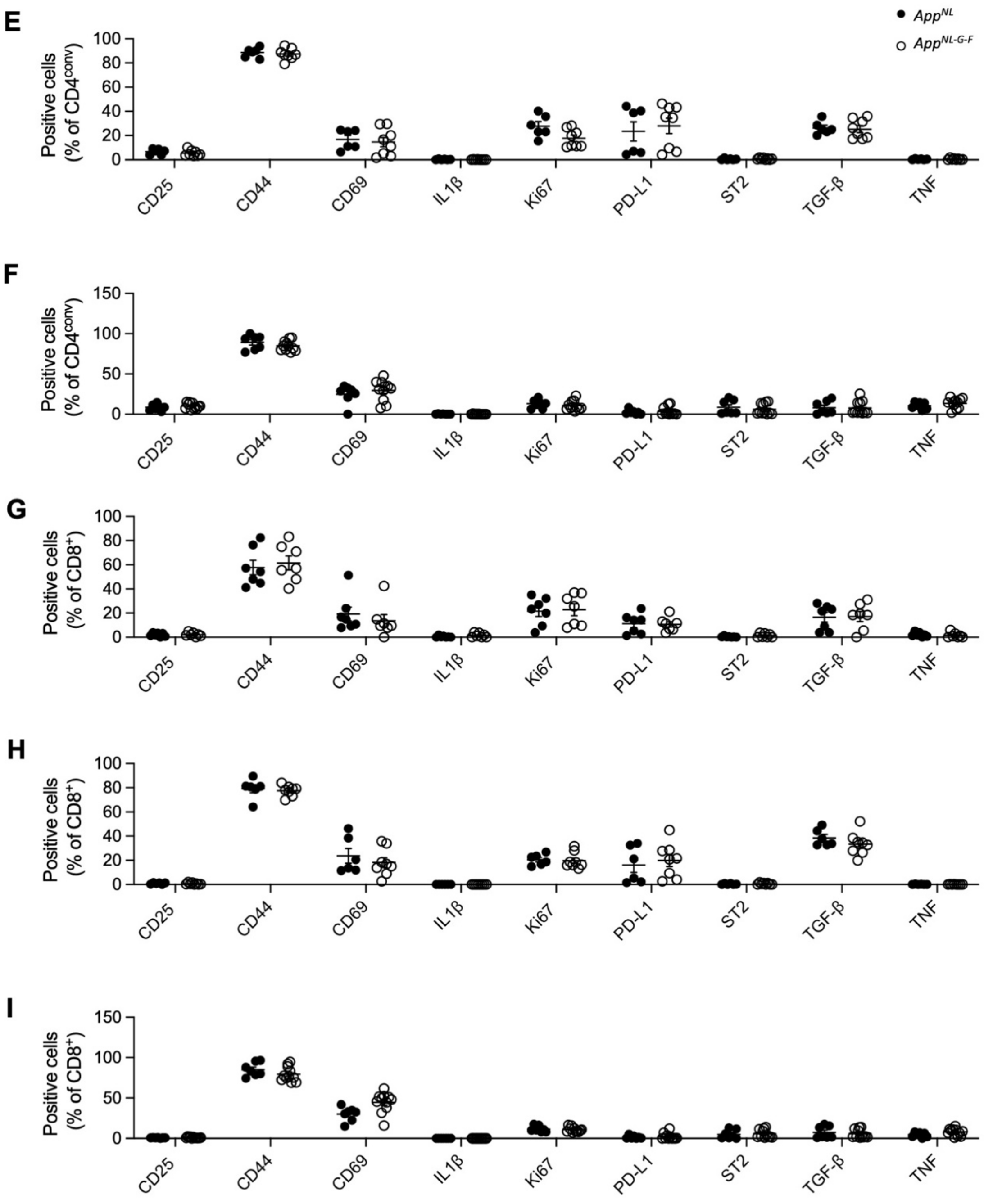

E
F
G
H
I
App^NL
App^NL-G-F
Positive cells (% of CD4conv)
Positive cells (% of CD4conv)
Positive cells (% of CD8+)
Positive cells (% of CD8+)
Positive cells (% of CD8+)
0
20
40
60
80
100
50
150
CD25
CD44
CD69
IL1β
Ki67
PD-L1
ST2
TGF-β
TNF

**Supplementary Figure 6. Flow cytometry analysis of peripheral CD4$^{conv}$ T cells in APP knock-in mice overexpressing IL2 (in the periphery?).** Perfused mice ($App^{NL}$, $App^{NL}$–IL2, $App^{NL\text{-}G\text{-}F}$ and $App^{NL\text{-}G\text{-}F}$–IL2 lines) were compared by high-dimensional flow cytometry for T cell immune profiles. Frequency of CD25, CD44$^{+}$CD62L$^{-}$, CD44$^{+}$CD62L$^{+}$, CD44$^{-}$CD62L$^{+}$, CD69, CD103, CTLA4, Helios, ICOS, Ki67, KLRG1, Neuropilin1, PD1 and ST2 expression within the peripheral CD4$^{conv}$ T cells in spleen at **A)** 2 (n=7,8,8,7), **B)** 4 (n=9,9,9,9) and **C)** 9 (n=9,10,10,7) months of age; in cLN at **D)** 2 (n=7,8,8,7), **E)** 4 (n=9,9,9,9) and **F)** 9 (n=9,10,10,7) months of age; and in the blood at **G)** 2 (n=7,8,8,7), **H)** 4 (n=9,9,9,9) and **I)** 9 (n=9,10,10,7) months of age.

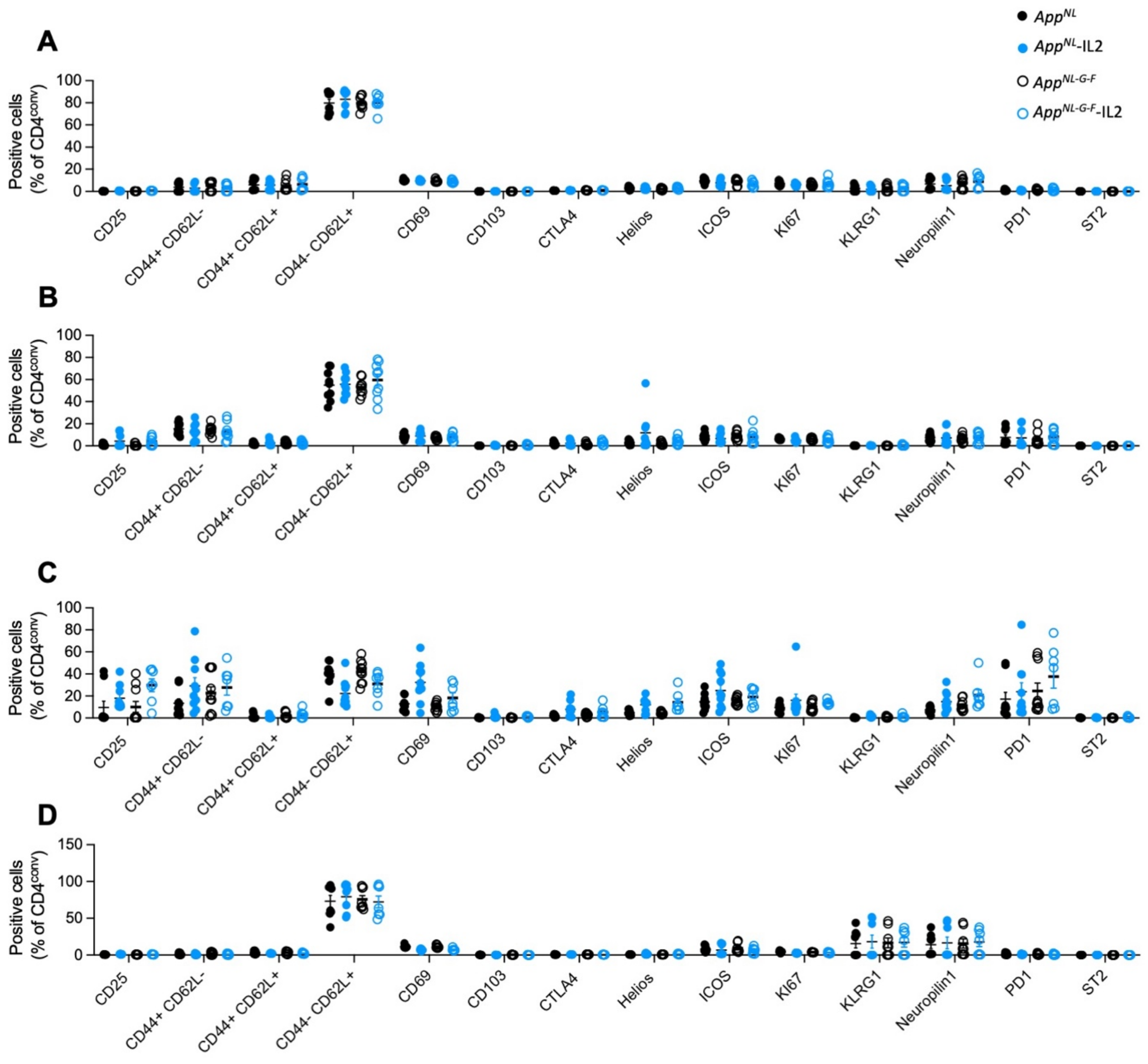

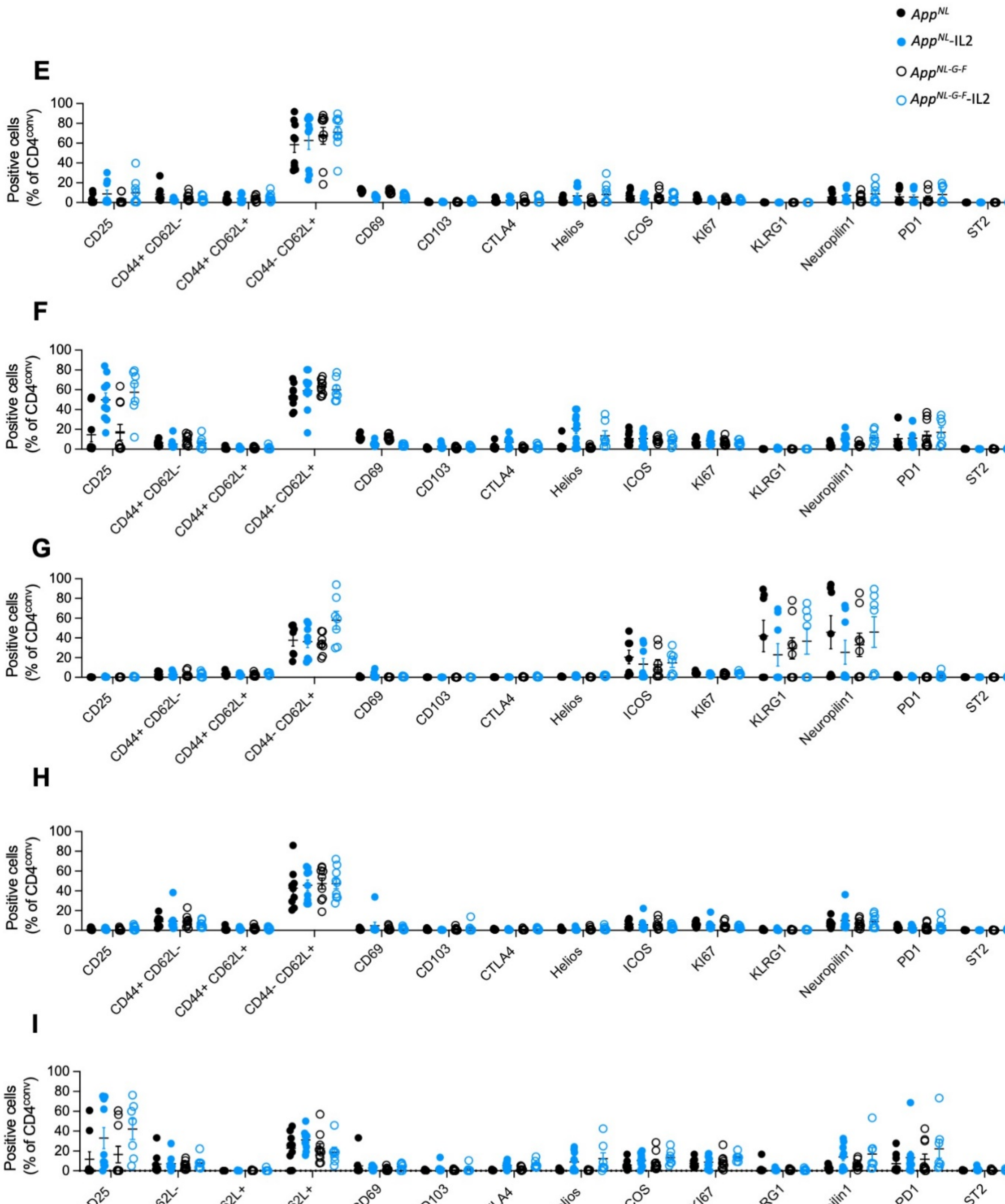

$App^{NL}$
$App^{NL}$-IL2
$App^{NL\text{-}G\text{-}F}$
$App^{NL\text{-}G\text{-}F}$-IL2
E
F
G
H
I
Positive cells (% of $CD4^{conv}$)
0
20
40
60
80
100
CD25
CD44+ CD62L-
CD44+ CD62L+
CD44- CD62L+
CD69
CD103
CTLA4
Helios
ICOS
KI67
KLRG1
Neuropilin1
PD1
ST2

**Supplementary Figure 7. Flow cytometry analysis of peripheral CD8 T cells in APP knock-in mice overexpressing IL2 (in the periphery?).** Perfused mice ( *App*$^{NL}$, *App*$^{NL}$–IL2, *App*$^{NL\text{-}G\text{-}F}$ and *App*$^{NL\text{-}G\text{-}F}$–IL2 lines) were compared by high-dimensional flow cytometry for T cell immune profiles. Frequency of CD25, CD44$^{+}$CD62L$^{-}$, CD44$^{+}$CD62L$^{+}$, CD44$^{-}$CD62L$^{+}$, CD69, CD103, CTLA4, Helios, ICOS, Ki67, KLRG1, Neuropilin1, PD1 and ST2 expression within the peripheral CD8 T cells in spleen at **A)** 2 (n=7,8,8,7), **B)** 4 (n=9,9,9,9) and **C)** 9 (n=9,10,10,7) months old; in cLN at **D)** 2 (n=7,8,8,7), **E)** 4 (n=9,9,9,9) and **F)** 9 (n=9,10,10,7) months old; and in the blood at **G)** 2 (n=7,8,8,7), **H)** 4 (n=9,9,9,9) and **I)** 9 (n=9,10,10,7) months old.

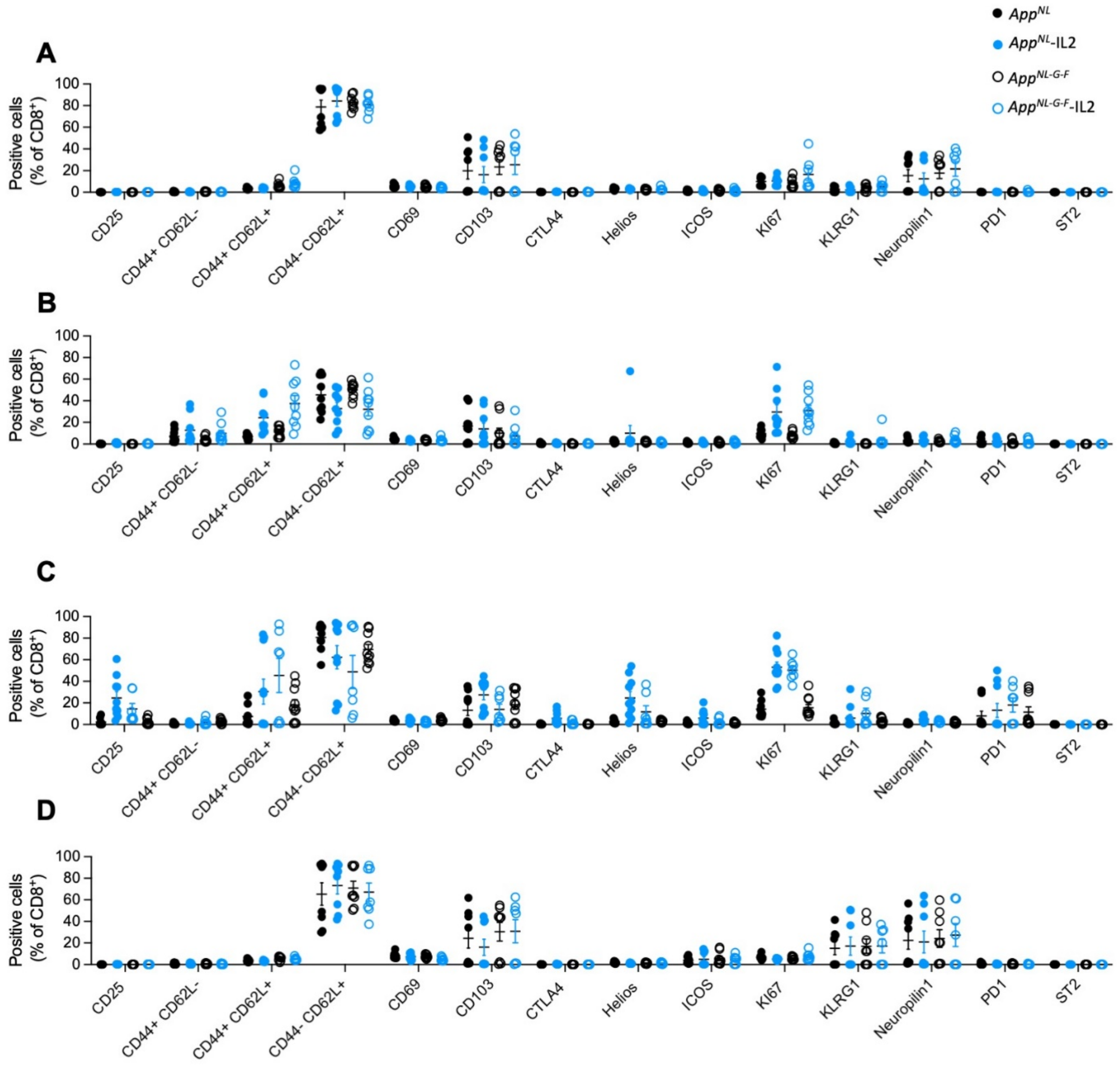

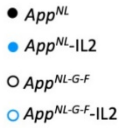

AppNL
AppNL-IL2
AppNL-G-F
AppNL-G-F-IL2


E

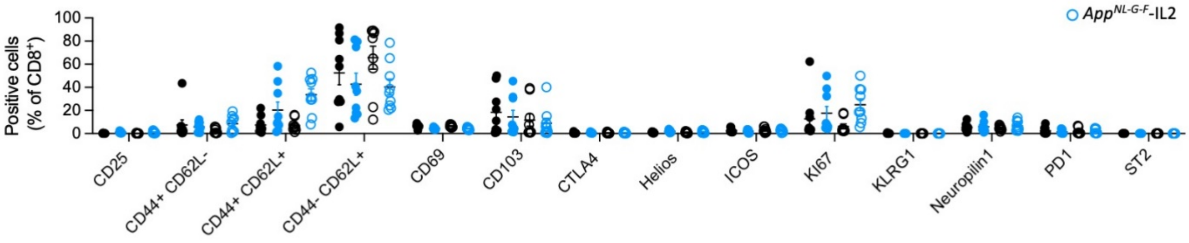

Positive cells (% of CD8+)
CD25
CD44+ CD62L-
CD44+ CD62L+
CD44- CD62L+
CD69
CD103
CTLA4
Helios
ICOS
KI67
KLRG1
Neuropilin1
PD1
ST2


F

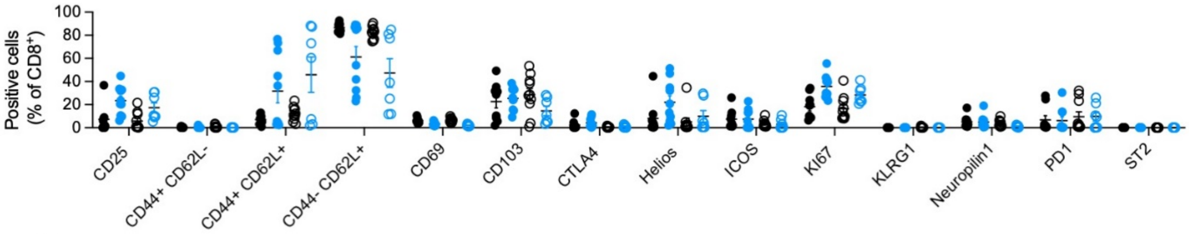

Positive cells (% of CD8+)
CD25
CD44+ CD62L-
CD44+ CD62L+
CD44- CD62L+
CD69
CD103
CTLA4
Helios
ICOS
KI67
KLRG1
Neuropilin1
PD1
ST2


G

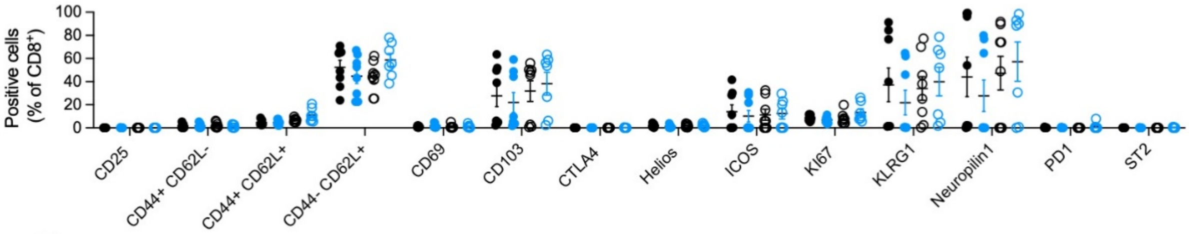

Positive cells (% of CD8+)
CD25
CD44+ CD62L-
CD44+ CD62L+
CD44- CD62L+
CD69
CD103
CTLA4
Helios
ICOS
KI67
KLRG1
Neuropilin1
PD1
ST2


H

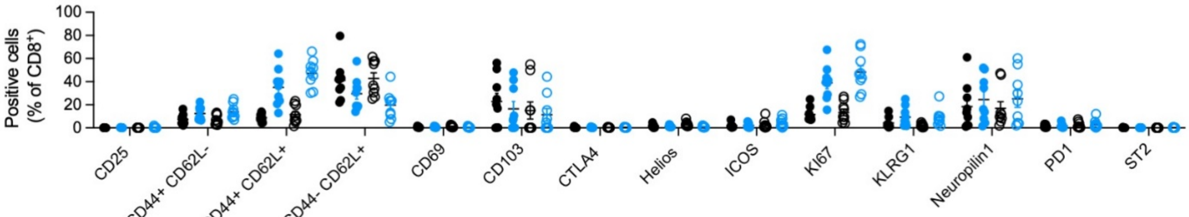

Positive cells (% of CD8+)
CD25
CD44+ CD62L-
CD44+ CD62L+
CD44- CD62L+
CD69
CD103
CTLA4
Helios
ICOS
KI67
KLRG1
Neuropilin1
PD1
ST2


I

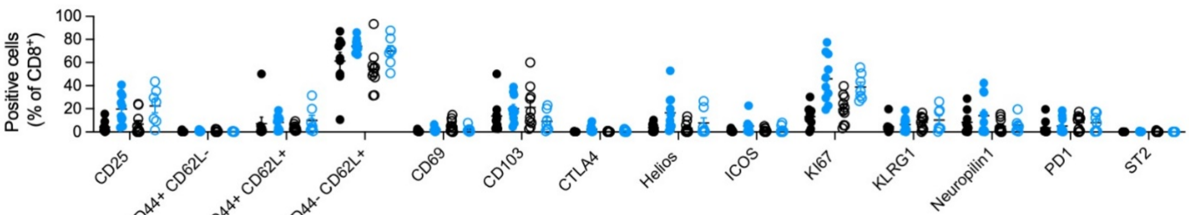

Positive cells (% of CD8+)
CD25
CD44+ CD62L-
CD44+ CD62L+
CD44- CD62L+
CD69
CD103
CTLA4
Helios
ICOS
KI67
KLRG1
Neuropilin1
PD1
ST2

**Supplementary Figure 8. Flow cytometry analysis of T cells in APP knock-in mice overexpressing IL2 (in the periphery?).** Perfused mice ( *App*$^{NL}$, *App*$^{NL}$–IL2, *App*$^{NL\text{-}G\text{-}F}$ and *App*$^{NL\text{-}G\text{-}F}$–IL2 lines) were compared by high-dimensional flow cytometry to establish the immune profile of the brain T cell population. Frequency of CD25, CD44, CD69, IL-1ß, Ki67, PD-L1, ST2, TGF-ß, and TNF expression in brain Tregs at **A)** 2 (n=7,8,8,7), **B)** 4 (n=9,9,9,9) and **C)** 9 (n=7,8,6,7) months of age; CD4$^{conv}$ T cells at **D)** 2 (n=7,8,8,7), **E)** 4 (n=9,9,9,9) and **F)** 9 (n=7,8,6,7) months of age; and within brain CD8 T cells at **G)** 2 (n=7,8,8,7), **H)** 4 (n=9,9,9,9) and **I)** 9 (n=7,8,6,7) months of age.

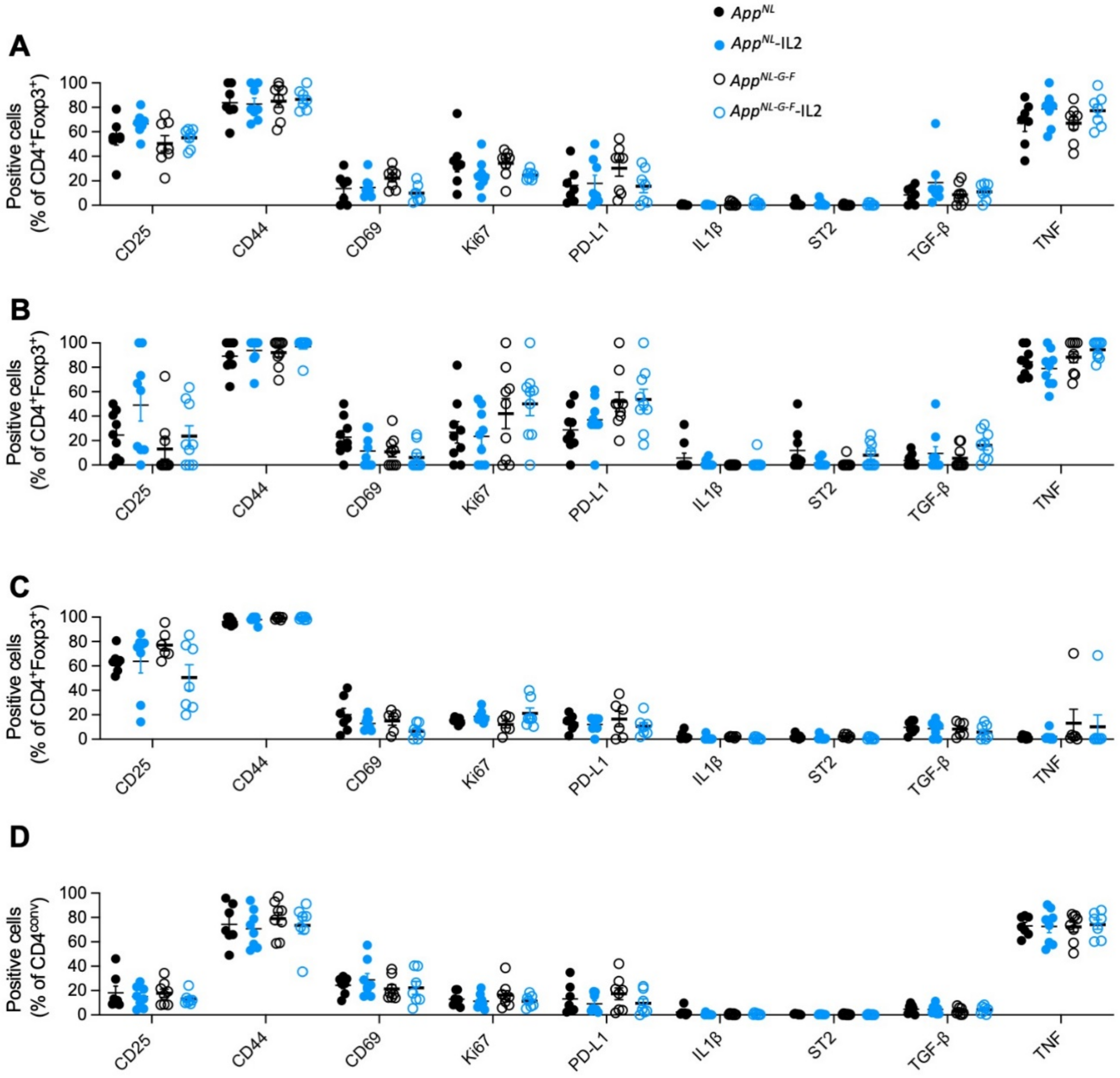

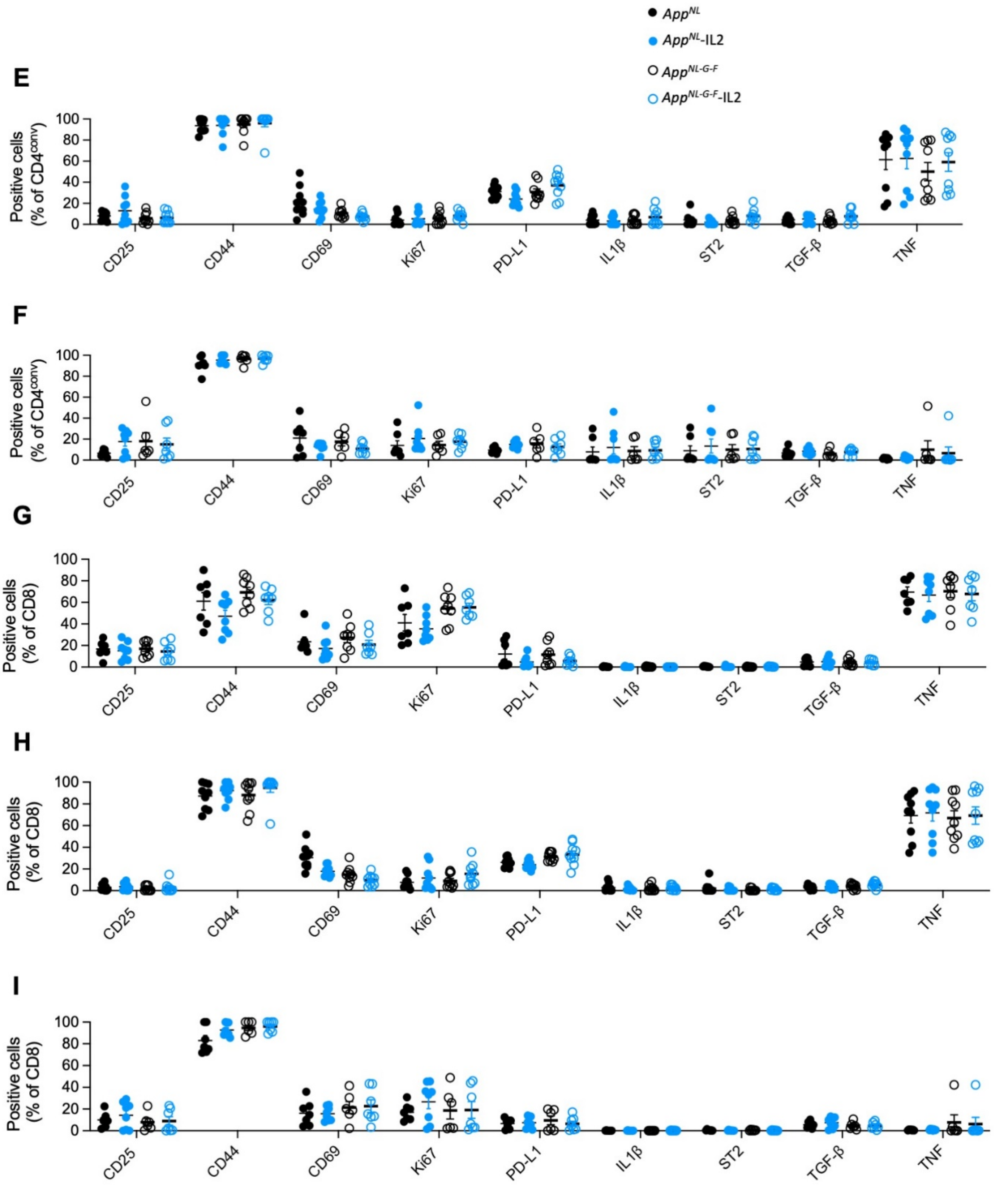


**Supplementary Figure 9. Flow cytometry analysis of peripheral Tregs in APP$^{NLGF}$-IL2 mice.** Perfused mice (APP$^{NL}$, APP$^{NL}$–IL2, APP$^{NLGF}$ and APP$^{NLGF}$–IL2 lines) were compared by high-dimensional flow cytometry to obtain the immune profiles present within the T cell population. Frequency of CD25, CD44$^+$CD62L$^-$, CD44$^+$CD62L$^+$, CD44$^-$CD62L$^+$, CD69, CD103, CTLA4, Helios, ICOS, Ki67, KLRG1, Neuropilin1, PD1 and ST2 expression within the peripheral Tregs in spleen at **A)** 2 (n=7,8,8,7), **B)** 4 (n=9,9,9,9) and **C)** 9 (n=9,10,10,7) months of age; in cLN at **D)** 2 (n=7,8,8,7), **E)** 4 (n=9,9,9,9) and **F)** 9 (n=9,10,10,7) months of age; and in the blood at **G)** 2 (n=7,8,8,7), **H)** 4 (n=9,9,9,9) and **I)** 9 (n=9,10,10,7) months of age.

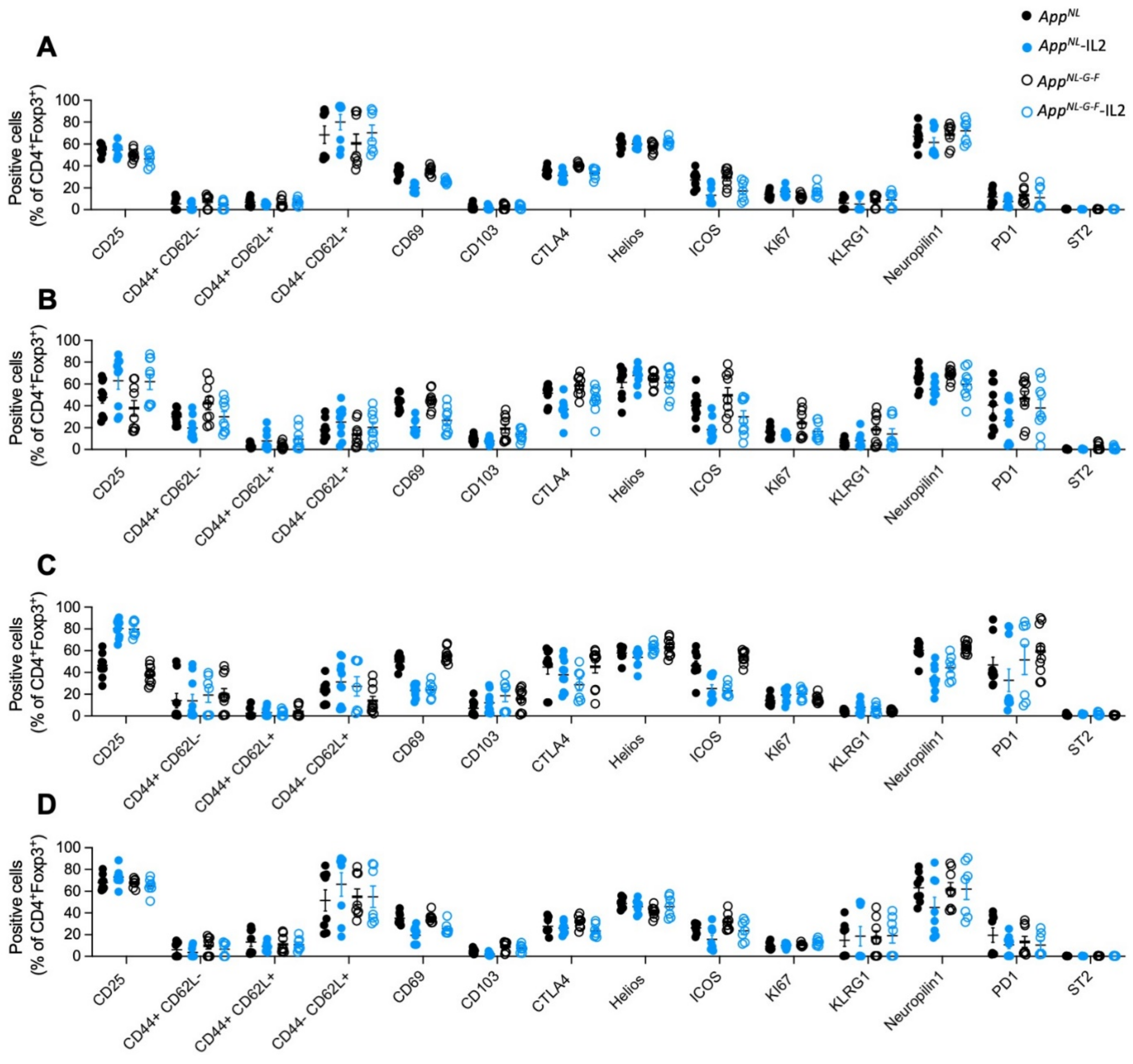

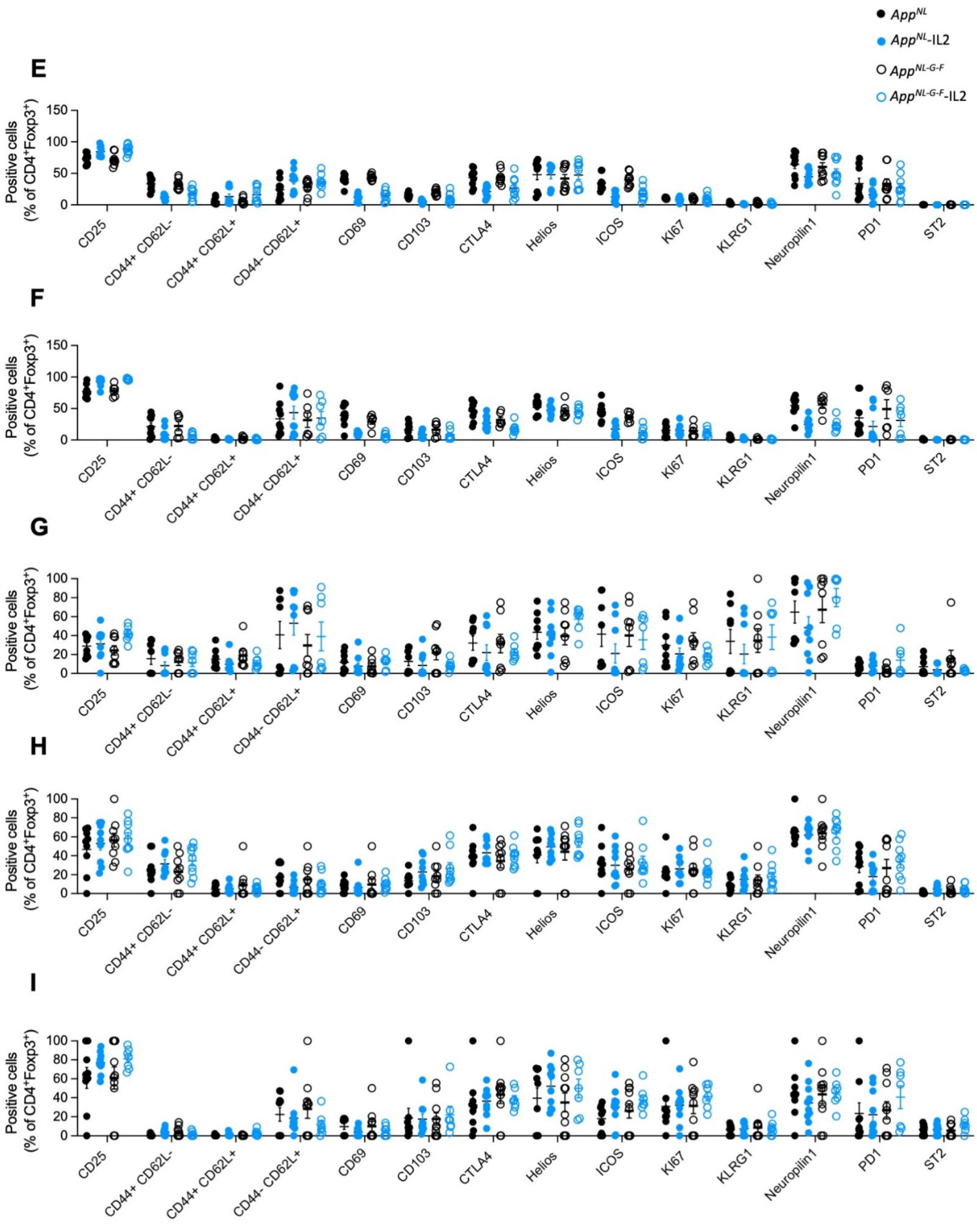

E
F
G
H
I
AppNL
AppNL-IL2
AppNL-G-F
AppNL-G-F-IL2
Positive cells
(% of CD4+Foxp3+)
CD25
CD44+ CD62L-
CD44+ CD62L+
CD44- CD62L+
CD69
CD103
CTLA4
Helios
ICOS
KI67
KLRG1
Neuropilin1
PD1
ST2

**Supplementary Figure 10. Flow cytometry analysis of peripheral CD4$^{conv}$ T cells in APP knock-in mice overexpressing IL2 in the CNS.** Perfused mice (*App*$^{NL\text{-}G\text{-}F}$ and *App*$^{NL\text{-}G\text{-}F}$–CamK2$^{IL2}$ lines) were compared by high-dimensional flow cytometry to obtain the immune profiles in the T cell population. Frequency of CD25, CD44$^{+}$CD62L$^{-}$, CD44$^{+}$CD62L$^{+}$, CD44$^{-}$CD62L$^{+}$, CD69, CD103, CTLA4, Helios, ICOS, Ki67, KLRG1, Neuropilin1, PD1 and ST2 expression within the peripheral CD4$^{conv}$ populations found in spleen at **A)** 2 (n=7,8), **B)** 4 (n=7,8) and **C)** 9 (n=8,8) months of age; in cLN at **D)** 2 (n=7,8), **E)** 4 (n=7,8) and **F)** 9 (n=8,8) months of age; and in the blood at **G)** 2 (n=7,8), **H)** 4 (n=7,8) and **I)** 9 (n=8,8) months of age.

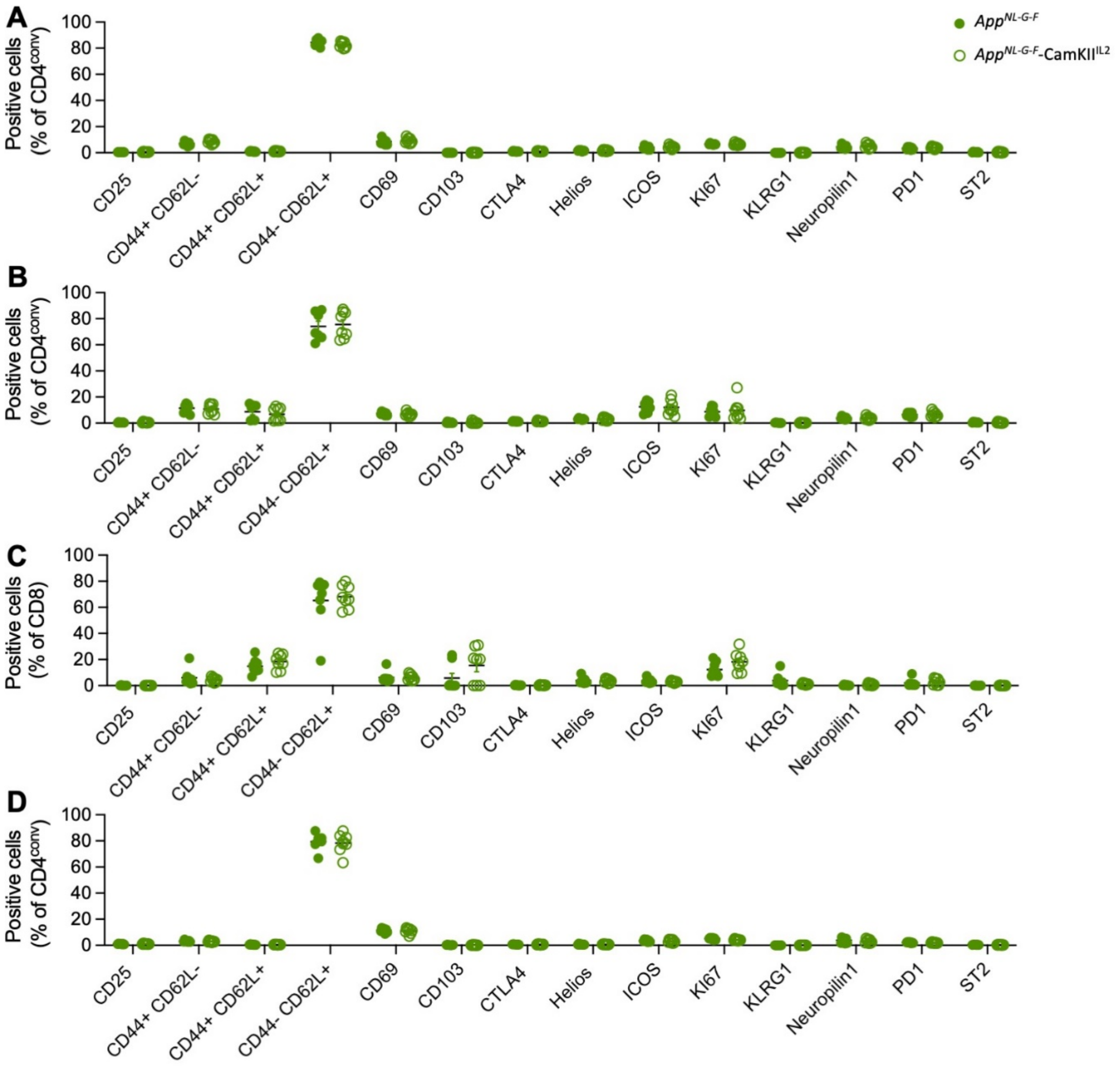

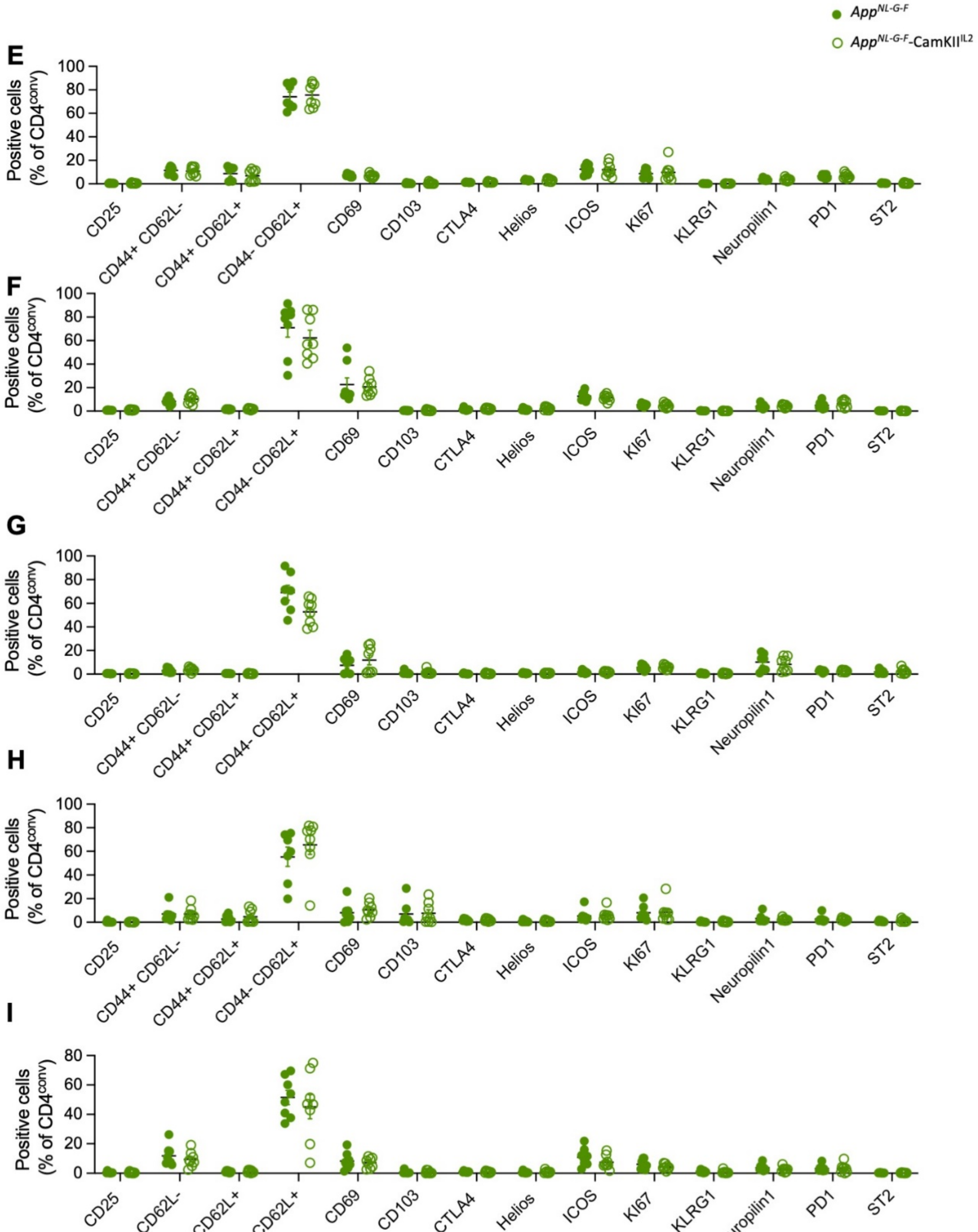

App^NL-G-F
App^NL-G-F-CamKII^IL2
E
F
G
H
I
Positive cells (% of CD4^conv)
0
20
40
60
80
100
CD25
CD44+ CD62L-
CD44+ CD62L+
CD44- CD62L+
CD69
CD103
CTLA4
Helios
ICOS
KI67
KLRG1
Neuropilin1
PD1
ST2

**Supplementary Figure 11. Flow cytometry analysis of peripheral CD8 T cells in APP knock-in mice overexpressing IL2 in the CNS.** Perfused mice ( $App^{NL\text{-}G\text{-}F}$ and $App^{NL\text{-}G\text{-}F}$–CamK2$^{IL2}$ lines) were compared by high-dimensional flow cytometry to obtain the immune profiles in the T cell population. Frequency of CD25, CD44$^+$CD62L$^-$, CD44$^+$CD62L$^+$, CD44$^-$CD62L$^+$, CD69, CD103, CTLA4, Helios, ICOS, Ki67, KLRG1, Neuropilin1, PD1 and ST2 expression within the peripheral CD8 T cell populations in spleen at **A)** 2 (n=7,8), **B)** 4 (n=7,8) and **C)** 9 (n=8,8) months of age; in cLN at **D)** 2 (n=7,8), **E)** 4 (n=7,8) and **F)** 9 (n=8,8) months of age; and in the blood at **G)** 2 (n=7,8), **H)** 4 (n=7,8) and **I)** 9 (n=8,8) months of age.

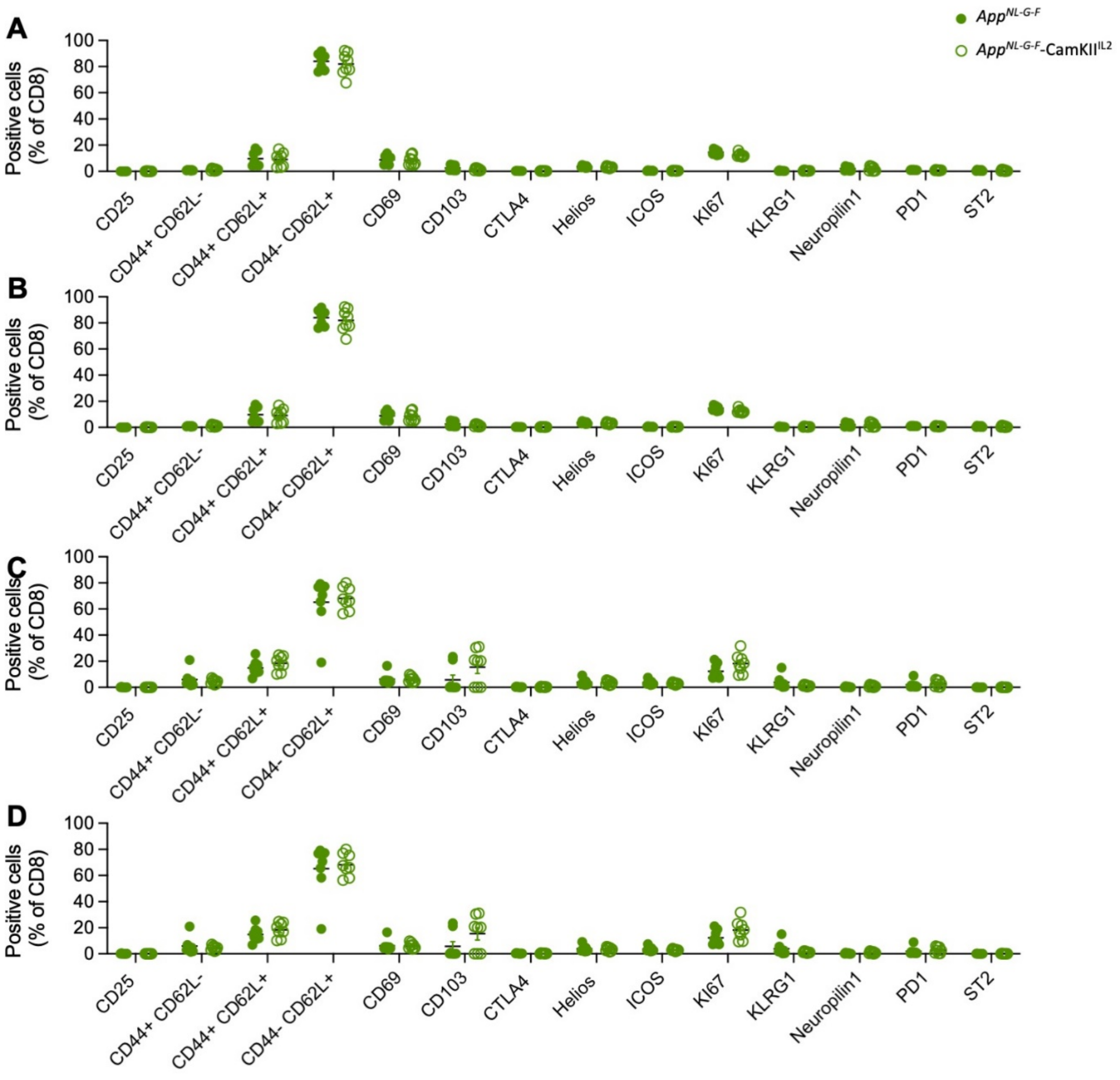

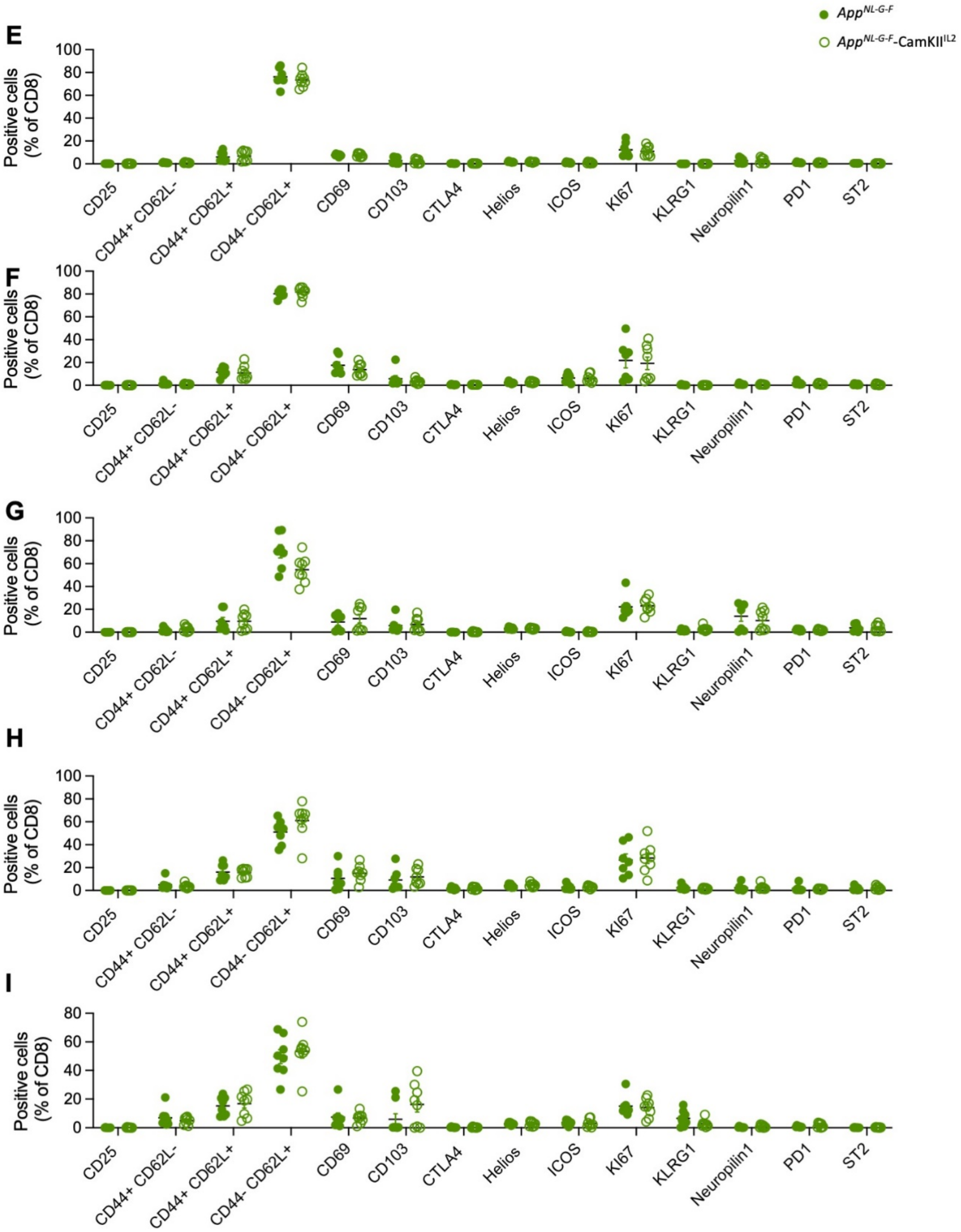

App^NL-G-F
App^NL-G-F-CamKII^IL2
E
Positive cells (% of CD8)
100
80
60
40
20
0
CD25
CD44+ CD62L-
CD44+ CD62L+
CD44- CD62L+
CD69
CD103
CTLA4
Helios
ICOS
KI67
KLRG1
Neuropilin1
PD1
ST2
F
Positive cells (% of CD8)
100
80
60
40
20
0
CD25
CD44+ CD62L-
CD44+ CD62L+
CD44- CD62L+
CD69
CD103
CTLA4
Helios
ICOS
KI67
KLRG1
Neuropilin1
PD1
ST2
G
Positive cells (% of CD8)
100
80
60
40
20
0
CD25
CD44+ CD62L-
CD44+ CD62L+
CD44- CD62L+
CD69
CD103
CTLA4
Helios
ICOS
KI67
KLRG1
Neuropilin1
PD1
ST2
H
Positive cells (% of CD8)
100
80
60
40
20
0
CD25
CD44+ CD62L-
CD44+ CD62L+
CD44- CD62L+
CD69
CD103
CTLA4
Helios
ICOS
KI67
KLRG1
Neuropilin1
PD1
ST2
I
Positive cells (% of CD8)
80
60
40
20
0
CD25
CD44+ CD62L-
CD44+ CD62L+
CD44- CD62L+
CD69
CD103
CTLA4
Helios
ICOS
KI67
KLRG1
Neuropilin1
PD1
ST2

**Supplementary Figure 12. Flow cytometry analysis of brain T cells in APP knock-in mice overexpressing IL2 in the CNS.** Perfused mice (*App*$^{NL\text{-}G\text{-}F}$ and *App*$^{NL\text{-}G\text{-}F}$–CamK2$^{IL2}$ lines) were compared by high-dimensional flow cytometry to obtain brain T cell immune profiles. Frequency of CD25, CD44, CD69, IL-1ß, Ki67, PD-L1, ST2, TGF-ß, and TNF expression within brain Tregs at **A)** 2 (n=7,8), **B)** 4 (n=9,9) and **C)** 9 (n=8,8) months of age; CD4$^{conv}$ T cells at **D)** 2 (n=7,8), **E)** 4 (n=9,9) and **F)** 9 (n=8,8) months of age; and CD8 T cells at **G)** 2 (n=7,8), **H)** 4 (n=9,9) and **I)** 9 (n=8,8) months of age.

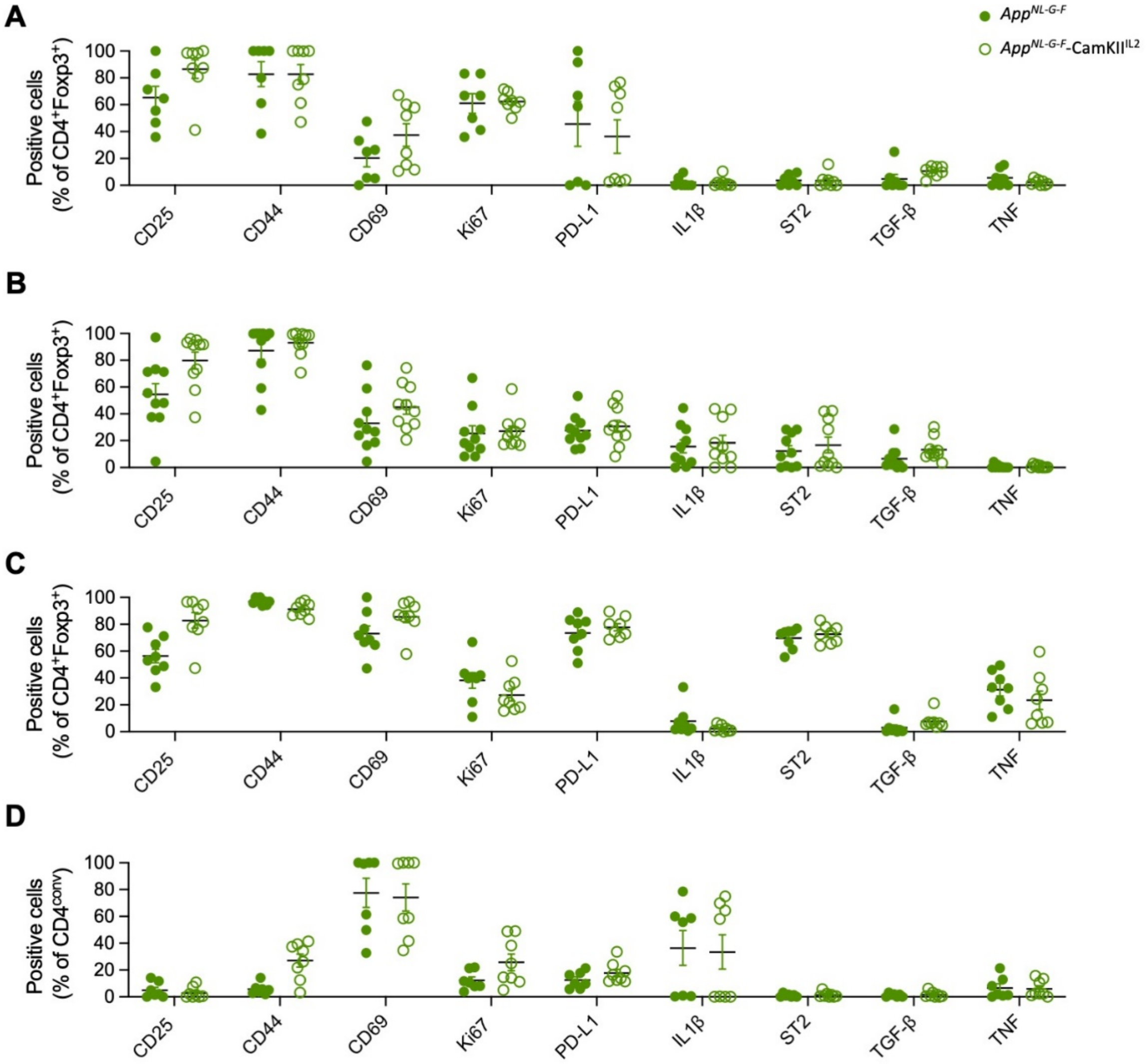

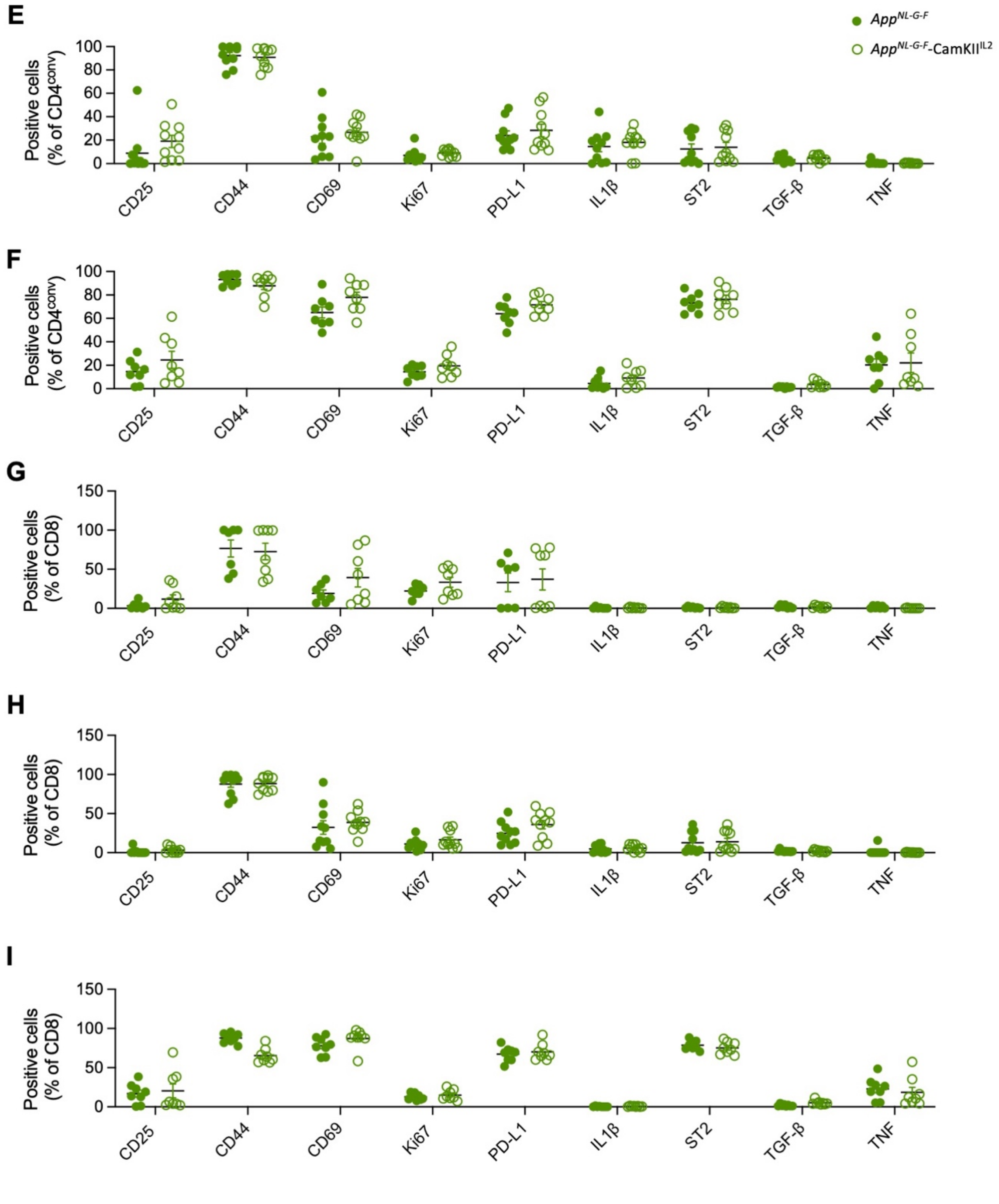
E
F
G
H
I
App^NL-G-F
App^NL-G-F-CamKII^IL2
Positive cells (% of CD4^conv)
Positive cells (% of CD8)
CD25
CD44
CD69
Ki67
PD-L1
IL1β
ST2
TGF-β
TNF

**Supplementary Figure 13. Flow cytometry analysis of peripheral Tregs in APP knock-in mice overexpressing IL2 in the CNS.** Perfused mice ( $App^{NL\text{-}G\text{-}F}$ and $App^{NL\text{-}G\text{-}F}$–CamK2$^{IL2}$ lines) were compared using high-dimensional flow cytometry to obtain T cell immune profiles. Frequency of CD25, CD44$^{+}$CD62L$^{-}$, CD44$^{+}$CD62L$^{+}$, CD44$^{-}$CD62L$^{+}$, CD69, CD103, CTLA4, Helios, ICOS, Ki67, KLRG1, Neuropilin1, PD1 and ST2 expression within the peripheral Tregs in spleen at **A)** 2 (n=7,8), **B)** 4 (n=7,8) and **C)** 9 (n=8,8) months of age; in cLN at **D)** 2 (n=7,8), **E)** 4 (n=7,8) and **F)** 9 (n=8,8) months of age; and in the blood at **G)** 2 (n=7,8), **H)** 4 (n=7,8) and **I)** 9 (n=8,8) months of age.

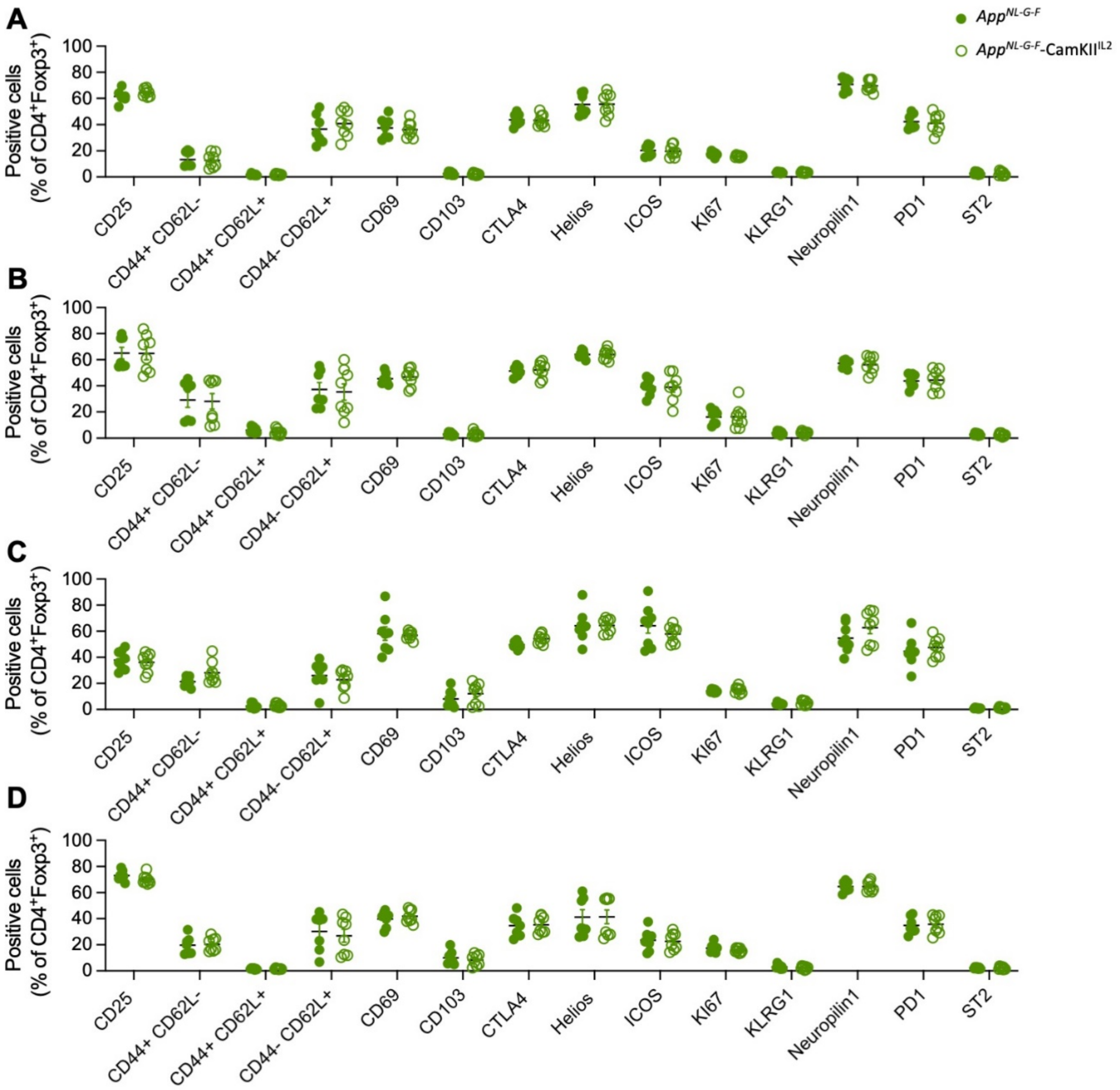

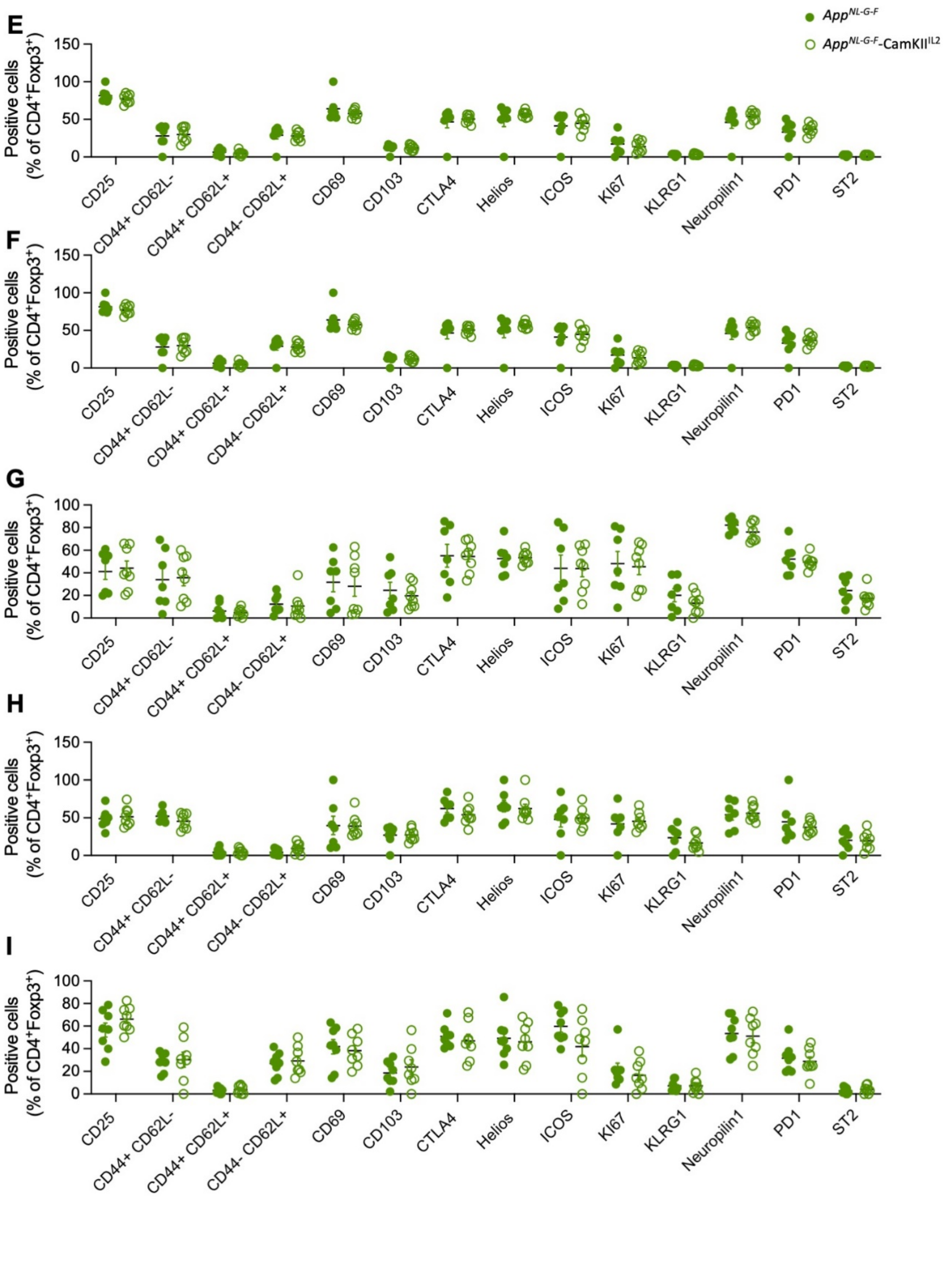
E
F
G
H
I
App^NL-G-F
App^NL-G-F-CamKII^IL2
Positive cells
(% of CD4+Foxp3+)
CD25
CD44+ CD62L-
CD44+ CD62L+
CD44- CD62L+
CD69
CD103
CTLA4
Helios
ICOS
KI67
KLRG1
Neuropilin1
PD1
ST2
